# Automated model discovery for skin: Discovering the best model, data, and experiment

**DOI:** 10.1101/2022.12.19.520979

**Authors:** Kevin Linka, Adrian Buganza Tepole, Gerhard A. Holzapfel, Ellen Kuhl

## Abstract

Choosing the best constitutive model and the right set of model parameters is at the heart of continuum mechanics. For decades, the gold standard in constitutive modeling has been to first select a model and then fit its parameters to data. However, the success of this approach is highly dependent on user experience and personal preference. Here we propose a new method that simultaneously and fully autonomously discovers the best model and parameters to explain experimental data. Mathematically, the model finding is translated into a complex non-convex optimization problem. We solve this problem by formulating it as a neural network, and leveraging the success, robustness, and stability of the optimization tools developed in classical neural network modeling. Instead of using a classical off-the-shelf neural network, we design a new family of Constitutive Artificial Neural Networks with activation functions that feature popular constitutive models and parameters that have a clear physical interpretation. Our new network inherently satisfies general kinematic, thermodynamic, and physical constraints and trains robustly, even with sparse data. We illustrate its potential for biaxial extension experiments on skin and demonstrate that the majority of network weights train to zero, while the small subset of non-zero weights defines the discovered model. Unlike classical network weights, these weights are physically interpretable and translate naturally into engineering parameters and microstructural features such as stiffness and fiber orientation. Our results suggest that Constitutive Artificial Neural Networks enable automated model, parameter, and experiment discovery and could initiate a paradigm shift in constitutive modeling, from user-defined to automated model selection and parameterization. Our source code, data, and examples are available at https://github.com/LivingMatterLab/CANN.

## 1 Motivation

Neural networks are gaining increased popularity in the computational mechanics community and are increasingly used as function approximators in constitutive modeling [1]. Neural networks learn functions from data by minimizing a loss function [38]. In constitutive modeling, the function is a model for the stress, the data are measured stress-strain pairs, and the loss function is the mean squared error between model and data [14]. Classical neural networks have evolved into a powerful technology to interpolate or describe big data; however, they cannot extrapolate or predict beyond their training regime [35]. They are an excellent choice when we have no information about the underlying data, but they totally ignore our expert knowledge, anyone can actually violate kinematic, thermodynamic, or physical constraints [43]. In the materials physics community, this has raised the question of how best to integrate neural networks and constitutive modeling and ideally combine the best features of both [62].

Two successful but fundamentally different strategies have emerged to integrate physical knowledge into network models: Physics-Informed Neural Networks that add physical equations as additional constraints to the loss function [20]; and Constitutive Artificial Neural Networks that explicitly modify the network input, output, and architecture to tightly incorporate physical constraints into the network design [31]. The former is broadly applicable to any type of ordinary [32] or partial [44] differential equations, while the latter is specifically tailored to constitutive equations [31]. In fact, almost two decades ago, the first constitutive neural network with strain invariants as input, free-energy functions as output, and a single hidden layer with logistic activation functions in between was proposed for rubber-like materials [48]. It has recently regained attention in the constitutive modeling community [63], with applications for planing rubber sheets [31], sheets with holes [54], entire tires [48], parachute deployment [2], and plastic surgery [51]. An inherent limitation of all these success stories is that their neural network parameters do not have a clear intuitive interpretation and teach us little to nothing about the underlying physics [21]. This raises the question of whether and how we can use our expertise in constitutive modeling to design a new family of Constitutive Artificial Neural Networks that not only approximate stresses from data [14], but rather discover the best constitutive model and meaningful model parameters to explain experimental data. Here, we will prototype this idea for transversely isotropic composite materials with experimental data from skin.

The skin is the largest organ in our body and our interface to the outside world. Its stiffness is tightly regulated to protect the underlying tissues from environmental insults while allowing for movement and interaction with objects ranging from clothes to medical prostheses. The first biomechanical study of skin dates back to 1861, when the Austrian anatomist Karl Langer punctured circular holes in the human cadaver skin and discovered its anisotropy as the circular punches turned into ellipsoidal shapes [22]. This experiment produced the classic Langer lines, topological lines parallel to the natural orientation of collagen fibers in the dermis that have important implications in plastic and reconstructive surgery [8, 61]. However, it was not until more than 70 years later that scientists fully characterized the three-dimensional response of the skin using a custom-designed biaxial extension system [23]. These experiments revealed a transversely isotropic behavior with a stiff response parallel to Langer’s lines and a soft response perpendicular to it [24] and the characteristic stretch-stiffening [25] that we now commonly associate with soft collagenous tissues [18]. Based on these pioneering experiments, different experimental protocols have been proposed [19, 29, 30, 59] for probing the skin under uni-aixal tension [41], biaxial extension [24], or torsion [5]. A legitimate question, particularly with regard to constitutive modeling, is which experiment is best suited to identifying its parameters [27], or, in view of model finding, which experiment provides the richest data for training neural networks for the skin [34]. Our intuition suggests that uniaxial tension or strip tests parallel and perpendicular to Langer’s lines should provide the best insight into the constitutive behavior of the skin [41]. But is this really true, and if so, how can we formally quantify it?

In parallel to the numerous experiments to characterize the nonlinear transversely isotropic response of skin, a long list of constitutive models has been developed for this important tissue throughout the past fifty years. Yet, there is no definitive choice of constitutive equation that is best suited for a particular dataset, no universal model that can be used for different animal species, and no standard testing protocol that can guarantee robust model training. The general idea of this manuscript is to prototype a new method to autonomously discover the best model, parameters, and experiment to characterize the constitutive behavior of the skin. For this purpose, we revisit the basics of constitutive modeling in Section 2 and demonstrate in Section 3 how this expert knowledge can be integrated into a new family of Constitutive Artificial Neural Networks. In the Section 4, we briefly review the homogeneous deformation mode of biaxial extension and introduce the data we use to train our model in Section 5. We discuss our results, limitations, and future directions in Section 6 and close with a brief conclusion in Section 7.

## 2 Constitutive modeling

To characterize the deformation of a test sample, we introduce the deformation map ***φ*** that maps material particles **X** from the undeformed configuration to particles, ***x* = *φ*(*X*)**, in the deformed configuration [3, 18]. The gradient of the deformation map ***φ*** with respect to the undeformed coordinates **X** defines the deformation gradient ***F*** with the Jacobian *J*,

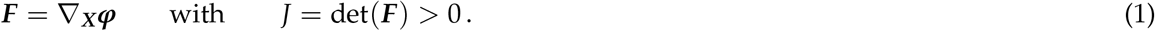

Multiplying ***F*** with its transpose ***F***^t^ from the left or the right introduces the right and left Cauchy–Green tensors **C** = ***F***^t^ · ***F*** and ***b*** = ***F*** · ***F***^t^, respectively. In the undeformed state, all three tensors are identical to the unit tensor, ***F*** = ***I***, **C** = ***I***, and ***b*** = ***I***, and the Jacobian is one, *J* = 1. A Jacobian smaller than one, 0 < *J* < 1, denotes compression and a Jacobian larger than one, 1 < *J*, denotes extension.

A *transversely isotropic* material is characterized through the pronounced direction ***n***_0_ with unit length || ***n***_0_ || = 1 in the reference configuration, the pronounced direction ***n*** = ***F*** · ***n***_0_ in the deformed configuration, and the associated structure tensor ***N*** = ***n***_0_ ⊗ ***n***_0_. We characterize its deformation state through the three principal invariants *I*_1_, *I*_2_, *I*_3_, and two additional invariants *I*_4_ and *I*_5_ [49],

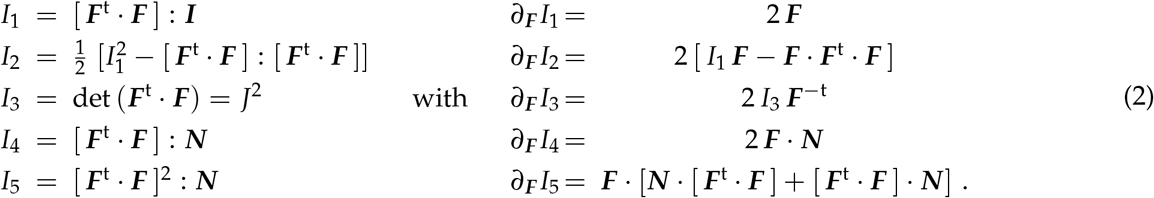

In the undeformed state, ***F*** = ***I***, and the five invariants are equal to three and one, *I*_1_ = 3, *I*_2_ = 3, *I*_3_ = 1, *I*_4_ = 1 and *I*_5_ = 1.

A *perfectly incompressible* material is characterized through a constant Jacobian equal to one, *I*_3_ = *J*^2^ = 1. Accordingly, its set of invariants reduces to four, *I*_1_, *I*_2_, *I*_4_, *I*_5_.

Next, we introduce the constitutive equation, a tensor-valued tensor function that defines the relation between the Piola or nominal stress ***P***, the force d***f*** per undeformed area d***A***, and the deformation gradient ***F*** [18, 56],

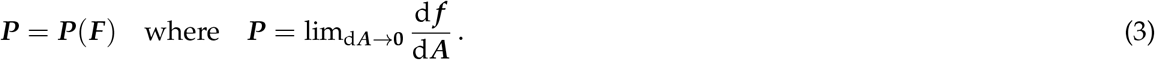

Here, instead of approximating the tensor-valued tensor function ***P***(***F***) directly through a classical neural network [14, 36, 47], our objective is to design a Constitutive Artificial Neural Network that limits the space of admissible functions by a priori guaranteeing common thermodynamic and physical constraints:

First, we consider *thermodynamic consistency* and imply that the Piola stress ***P*** follows from the second law of thermodynamics, 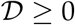 (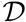 stands for dissipation), as the derivative of the free-energy function *ψ* with respect to the deformation gradient ***F*** [57],

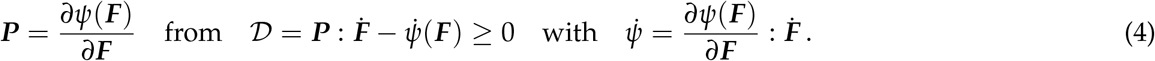

Second, we imply *material objectivity* [42] to ensure that our constitutive equation does not depend on the external frame of reference by requiring that the free energy *ψ* is a function of the right Cauchy-Green tensor, ***C*** = ***F***^t^ · ***F*** [56],

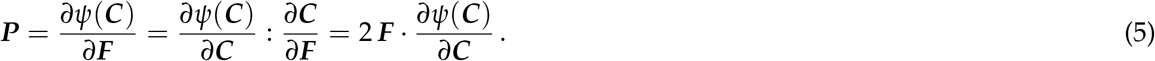

Third, we consider *material symmetry* for transversely isotropic materials by ensuring that the free energy *ψ* only depends on the five invariants from eq. (2) [18],

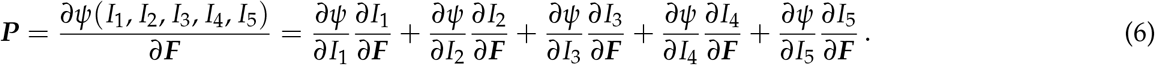

Fourth, we assume *perfect incompressibility* such that the third invariant remains constant, *I*_3_ = 1 = const., and we correct the free-energy function by a pressure term, –*p* ***F***^-t^, where 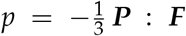 is the hydrostatic pressure, an additional unknown that we typically determine from the boundary conditions,

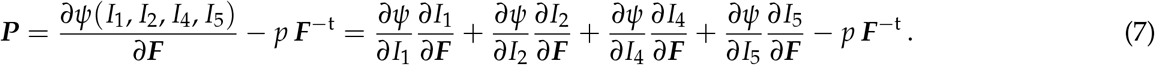

Fifth, we ensure *polyconvexity* [4] by considering a special subclass of free-energy functions *ψ*, which we can express as the sum of polyconvex subfunctions, *ψ*_1_, *ψ*_2_, *ψ*_4_, *ψ*_5_, of each invariant [12, 16], such that the stresses take the following additive form,

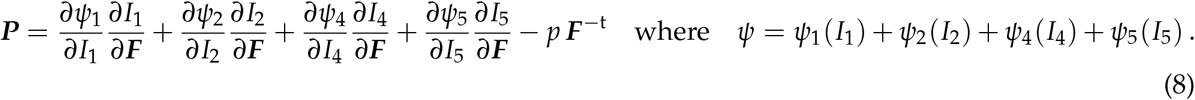

With the derivatives of the invariants from eq. (2), this results in the following explicit from,

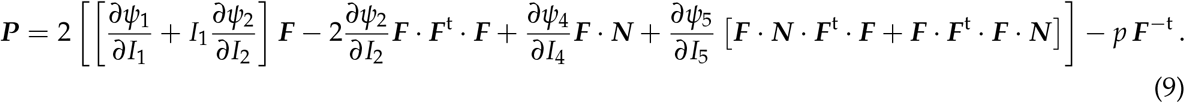

Finally, we consider physically reasonable constitutive restrictions [3], and ensure that the free energy *ψ* is *non-negative* for all deformation states, *ψ*(***F***) ≥ 0 for all ***F***; *zero* in the reference configuration, *ψ*(***F***) ≐ 0 at ***F*** = ***I***; and *infinite* for infinite compression or infinite expansion, *ψ*(***F***) → ∞ for *J* → 0 or *J* → ∞.

## 3 Constitutive Artificial Neural Networks

We now propose a new family of Constitutive Artificial Neural Networks that satisfy the conditions of thermodynamic consistency, material objectivity, material symmetry, perfect incompressibility, polyconvexity, and physical restrictions by design. Figure 1 illustrates an example of a transversely isotropic, perfectly incompressible Constitutive Artificial Neural Network with two hidden layers and four and eight nodes. The first layer generates powers (o) and (o)^2^ of the network input and the second layer applies the identity, (o) and the exponential function (exp(o)) to these powers. The constitutive equation of this networks takes the following explicit form, i.e.

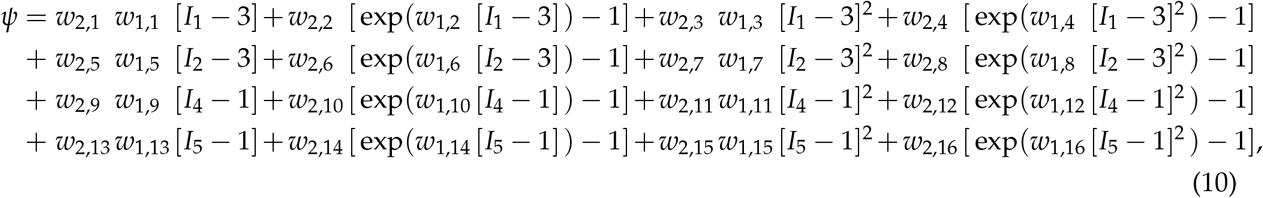

corrected by the pressure term *ψ* = *ψ* – *p* [*J* – 1]. Using the second law of thermodynamics, we can derive an explicit expression for the Piola stress, ***P*** = *∂ψ*/*∂***F**, i.e.

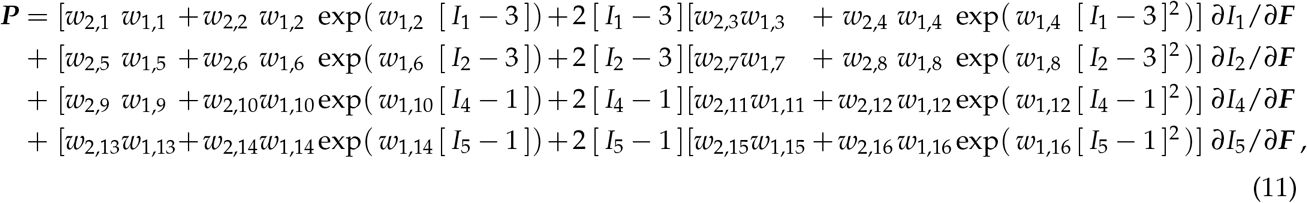

corrected by the pressure term *P* = *P* – *p* ***F***^-t^. For this particular format, one of the two weights with odd second indices becomes redundant, and we can reduce the set of network parameters from 32 to 24, ***w*** = [(*w*_1/1_*w*_2/1_), *w*_1/2_, *w*_2,2_, (*w*_1,3_*w*_2,3_), *w*_1,4_, *w*_2,4_, (*w*_1,5_*w*_2,5_), *w*_1,6_, *w*_2,6_, (*w*_1,7_*w*_2,7_), *w*_1,8_, *w*_2,8_, (*w*_1,9_*w*_2,9_), *w*_1,10_, *w*_2,10_, (*w*_1,11_*w*_2,11_), *w*_1,12_, *w*_2,12_, (*w*_1,13_ *w*_2,13_), *w*_1,14_, *w*_2,14_, (*w*_1,15_*w*_2,15_), *w*_1,16_, *w*_2,16_]. We learn the network weights **w** by minimizing a loss function *L* that penalizes the error between model and data. We characterize this error as the mean squared error, the *L*_2_-norm of the difference between the model ***P***(***F**_i_*) and data 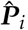, divided by the number of training points *n*_trn_,

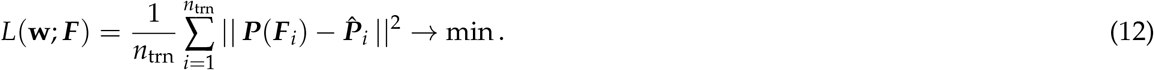

**Figure 1:**
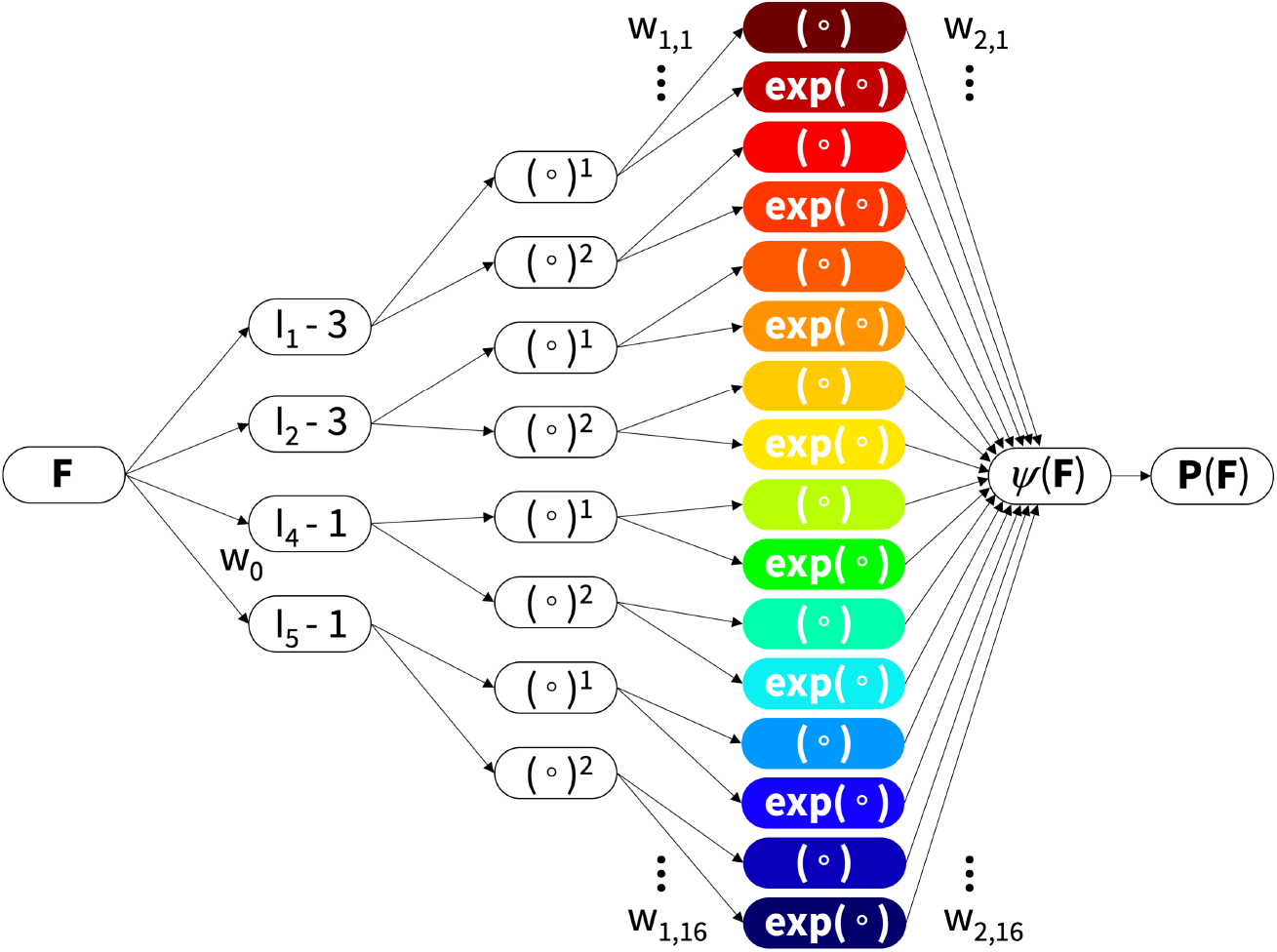
Constitutive Artificial Neural Network. Transversely isotropic, perfectly incompressible, Constitutive Artificial Neural Network with two hidden layers for approximating the single scalar-valued free-energy function *ψ*(*I*_1_, *I*_2_, *I*_4_, *I*_5_) as a function of the invariants of the deformation gradient ***F*** using sixteen terms. The first layer generates powers (o) and (o)^2^ of the network input and the second layer applies the identity (o) and exponential functions (exp(o)) to these powers. The networks are not fully connected by design to satisfy the condition of polyconvexity a priori.

We train the neural network by minimizing the loss function (12) and constraining the network weights to always remain non-negative, **w** ≥ **0**. Instead of implementing this minimization ourselves, we use the robustness and stability of the optimization tools developed for machine learning. In particular, we choose the widely used ADAM optimizer, a robust adaptive algorithm for gradient-based first-order optimization.

## 4 Biaxial extension tests

To discover the constitutive model for skin, we consider data from biaxial extension tests on rabbit [24, 25] and pig [50, 51] skin. We represent skin as a *transversally isotropic, perfectly incompressible* material using eq. (8),

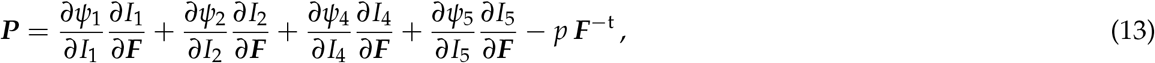

where *p* denotes the hydrostatic pressure, which we determine from the zero-thickness-stress condition. In *biaxial extension* tests, the skin specimen is stretched in two orthogonal directions, *λ*_1_ ≥ 1 and *λ*_2_ ≥ 1. From the incompressibility condition, 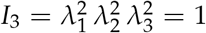, the stretch in the thickness direction, *λ*_3_ = (*λ*_1_ *λ*_2_)^−1^ ≤ 1, is uniquely defied through these two stretches. In all experiments, the samples are mounted with Langer’s lines along one of the stretch directions so that the deformation remains homogeneous and shear free, and the deformation gradient ***F*** and Piola stress ***P*** remain diagonal,

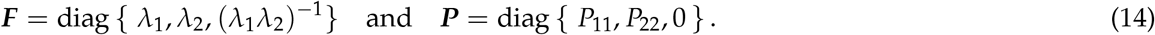

We use the explicit expressions of the invariants from eq. (2),

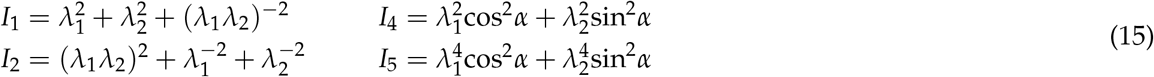

and their derivatives,

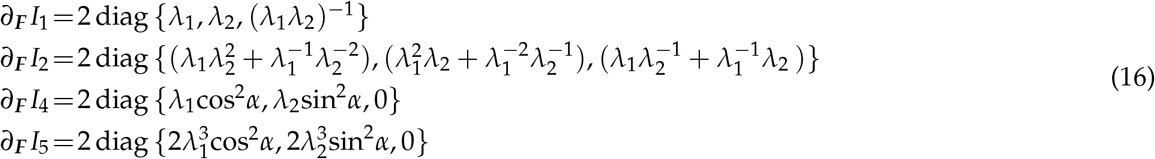

to determine the pressure *p* from the zero-thickness-stress condition in the third direction,

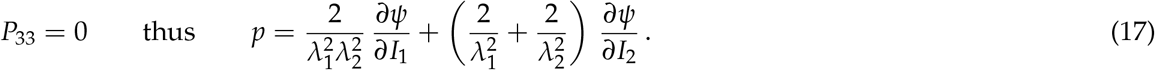

Equation (13) then provides explicit analytical expressions for the nominal stresses *P*_11_ and *P*_22_ in terms of the stretches *λ*_1_ and *λ*_2_,

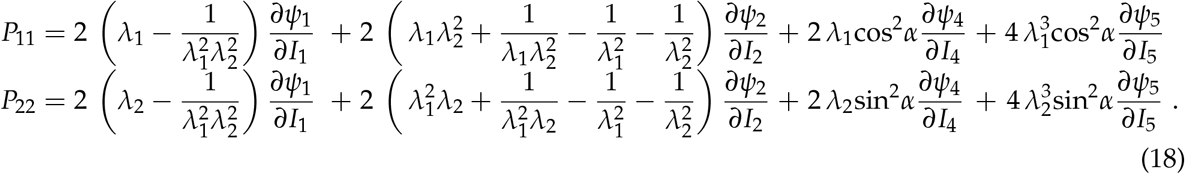

We compare the stress-stretch relations (18) with the very first reported biaxial extension experiments on the skin [24, 25] from almost half a century ago. These experiments include both strip-x and strip-y tests and biaxial extension tests on square rabbit skin samples with an average initial area of 35 × 35 mm^2^ and an average thickness of 1.2 mm. For the strip tests, the skin samples are either stretched in the *x*-direction, *λ*_1_ ≥ 1.000, with the *y*-direction fixed, *λ*_2_ = 1.000, or vice versa, resulting in two pairs of the individual datasets {*λ*_1_, *P*_11_} and {*λ*_2_, *P*_22_}. For the biaxial tests, the samples are stretched in the *x*-direction, *λ*_1_ ≥ 1.0, at four constant stretch levels in the *y*-direction, *λ*_2_ = 1.000, 1.087, 1.235, 1.415 =const., resulting in four triplets of datasets {*λ*_1_, *P*_11_, *P*_22_}. Table 1 summarizes discrete data pairs and triplets from the reported rabbit skin experiments [24, 25].

**Table 1:** Strip-x, strip-y, and biaxial extension data for rabbit skin. Skin samples are gradually stretched in one direction, while stretching is fixed in the orthogonal direction. The reported forces are converted in nominal stresses *P*_11_ and *P*_22_ for square 35 × 35 mm^2^ samples with a thickness of 1.2 mm [24, 25].

| rabbit strip-x |  | rabbit strip-y |  | rabbit biaxial |  |  | rabbit biaxial |  |  | rabbit biaxial |  |  | rabbit biaxial |  |  |
| --- | --- | --- | --- | --- | --- | --- | --- | --- | --- | --- | --- | --- | --- | --- | --- |
| $\lambda_2 = 1.000$ | | $\lambda_1 = 1.000$ | | $\lambda_2 = 1.000$ | | | $\lambda_2 = 1.087$ | | | $\lambda_2 = 1.235$ | | | $\lambda_2 = 1.415$ | | |
| $\lambda_1$<br>[-] | $P_{11}$<br>[kPa] | $\lambda_2$<br>[-] | $P_{22}$<br>[kPa] | $\lambda_1$<br>[-] | $P_{11}$<br>[kPa] | $P_{22}$<br>[kPa] | $\lambda_1$<br>[-] | $P_{11}$<br>[kPa] | $P_{22}$<br>[kPa] | $\lambda_1$<br>[-] | $P_{11}$<br>[kPa] | $P_{22}$<br>[kPa] | $\lambda_1$<br>[-] | $P_{11}$<br>[kPa] | $P_{22}$<br>[kPa] |
| 1.000 | 0.00 | 1.000 | 0.00 | 1.000 | 0.00 | 0.00 | 0.000 | 0.00 | 0.00 | 0.950 | 0.00 | 0.00 | 0.820 | 0.00 | 0.00 |
| 1.100 | 0.04 | 1.100 | 0.16 | 1.100 | 0.13 | 0.09 | 0.036 | 0.34 | 0.08 | 1.000 | 0.05 | 0.34 | 0.900 | 0.10 | 0.86 |
| 1.200 | 0.08 | 1.200 | 0.42 | 1.200 | 0.30 | 0.15 | 0.117 | 0.38 | 0.11 | 1.100 | 0.16 | 0.38 | 1.000 | 0.26 | 0.95 |
| 1.300 | 0.17 | 1.300 | 0.68 | 1.300 | 0.50 | 0.21 | 0.198 | 0.44 | 0.14 | 1.200 | 0.30 | 0.44 | 1.100 | 0.44 | 1.00 |
| 1.400 | 0.29 | 1.400 | 1.08 | 1.400 | 0.65 | 0.26 | 0.317 | 0.49 | 0.22 | 1.300 | 0.47 | 0.49 | 1.200 | 0.65 | 1.26 |
| 1.500 | 0.42 | 1.440 | 1.65 | 1.500 | 0.77 | 0.33 | 0.406 | 0.57 | 0.27 | 1.400 | 0.69 | 0.57 | 1.300 | 0.98 | 1.64 |
| 1.600 | 0.58 | 1.467 | 2.25 | 1.600 | 0.98 | 0.40 | 0.615 | 0.66 | 0.32 | 1.500 | 1.02 | 0.66 | 1.368 | 1.50 | 2.34 |
| 1.700 | 0.74 | 1.480 | 2.72 | 1.700 | 1.46 | 0.47 | 0.976 | 0.93 | 0.40 | 1.600 | 1.43 | 0.93 | 1.400 | 2.26 | 3.50 |
| 1.800 | 1.03 | 1.490 | 3.50 | 1.800 | 2.24 | 0.59 | 1.489 | 1.63 | 0.53 | 1.700 | 2.81 | 1.63 | 1.419 | 2.93 | 4.67 |
| 1.860 | 1.46 | 1.503 | 4.67 | 1.860 | 3.18 | 0.75 | 2.608 | 2.05 | 0.74 | 1.722 | 3.53 | 2.05 | 1.436 | 3.42 | 5.84 |
| 1.900 | 2.34 | 1.509 | 5.84 | 1.910 | 4.27 | 0.93 | 4.244 | 3.26 | 1.17 | 1.745 | 4.41 | 3.26 | 1.447 | 3.94 | 7.01 |
| 1.922 | 3.50 | 1.513 | 7.01 | 1.942 | 6.09 | 1.17 | 5.274 | 4.38 | 1.75 | 1.762 | 5.54 | 4.38 | 1.456 | 4.55 | 8.17 |
| 1.929 | 4.67 | 1.515 | 8.17 | 1.966 | 7.46 | 1.59 | 6.001 | 5.84 | 2.34 | 1.774 | 6.87 | 5.84 | 1.465 | 4.95 | 9.34 |
| 1.936 | 5.84 | 1.518 | 9.34 | 1.985 | 9.03 | 2.24 | 6.699 | 7.01 | 2.92 | 1.780 | 7.93 | 7.01 | 1.474 | 5.47 | 10.51 |
| 1.941 | 7.01 | 1.521 | 10.51 | 2.000 | 10.37 | 2.80 | 7.851 | 8.17 | 3.50 | 1.785 | 9.02 | 8.17 | 1.482 | 6.14 | 11.68 |
| 1.943 | 9.34 | 1.523 | 11.68 | 2.013 | 11.88 | 3.39 | 10.064 | 9.34 | 4.67 | 1.790 | 10.17 | 9.34 | 1.489 | 6.57 | 12.84 |
| 1.948 | 11.68 | 1.526 | 12.84 | 2.021 | 13.28 | 3.85 | 12.156 | 11.68 | 5.84 | 1.800 | 12.15 | 11.68 | 1.494 | 6.94 | 14.01 |
| 1.950 | 14.78 | 1.528 | 14.78 | 2.023 | 14.76 | 4.32 | 14.642 | 13.54 | 6.77 | 1.806 | 14.48 | 13.54 | 1.496 | 7.24 | 14.59 |

Next, we translate the nominal stresses (18) into the true stress *σ*_11_ and *σ*_22_, i.e.

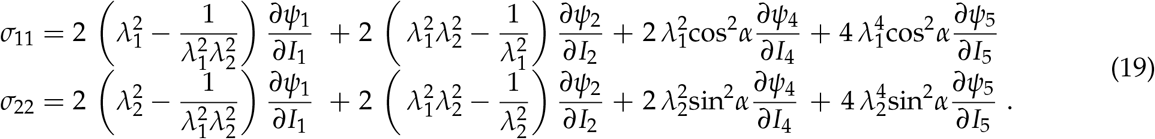

We compare the stress-stretch relations (19) with recent biaxial extension experiments on pig skin [50, 51]. These experiments include five sets of biaxial extension tests with prescribed stretch pairs, strip-x with *λ*_2_ = 1.000, off-x with 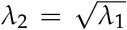, equi-biaxial with *λ*_2_ = *λ*_1_, off-y with 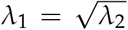, and strip-y with *λ*_1_ = 1.000. They provide five triplets of datasets, {*λ*_1_, *σ*_11_, *σ*_22_} or {*λ*_2_, *σ*_11_, *σ*_22_}, for which the second stretch *λ*_2_ or *λ*_1_ is either kept constant or increased as a function of *λ*_1_ or *λ*_2_, such that the maximum principal stretch is inclined at angles of 90.0^0^,67.5^0^,45.0^0^,22.5^0^,0.0^0^ towards the collagen fiber direction. Table 2 summarizes discrete data triplets from the pig skin experiments [50, 51].

**Table 2:** Biaxial extension data for pig skin. Skin samples are gradually stretched in two orthogonal directions. The ratio between the two stretches varies for all five experiments, such that the principal stretch axis is inclined by angles of 90.0^0^,67.5°,45.0^0^,22.5^0^,0.0^0^ in the collagen fiber orientation. Stresses are reported as true stresses *σ*_11_ and *σ*_22_ [50, 51].

| pig strip-x |  |  | pig off-x |  |  | pig biaxial |  |  | pig off-y |  |  | pig strip-y |  |  |
| --- | --- | --- | --- | --- | --- | --- | --- | --- | --- | --- | --- | --- | --- | --- |
| $\lambda_2 = 1.000$ | | | $\lambda_2 = \sqrt{\lambda_1}$ | | | $\lambda_1 = \lambda_2$ | | | $\lambda_1 = \sqrt{\lambda_2}$ | | | $\lambda_1 = 1.000$ | | |
| $\lambda_1$<br>[-] | $\sigma_{11}$<br>[MPa] | $\sigma_2$<br>[MPa] | $\lambda_1$<br>[-] | $\sigma_{11}$<br>[MPa] | $\sigma_2$<br>[MPa] | $\lambda_1, \lambda_2$<br>[-] | $\sigma_{11}$<br>[MPa] | $\sigma_2$<br>[MPa] | $\lambda_2$<br>[-] | $\sigma_{11}$<br>[MPa] | $\sigma_2$<br>[MPa] | $\lambda_2$<br>[-] | $\sigma_{11}$<br>[MPa] | $\sigma_2$<br>[MPa] |
| 1.000 | 0.000 | 0.000 | 1.000 | 0.000 | 0.000 | 1.000 | 0.000 | 0.000 | 1.000 | 0.000 | 0.000 | 1.000 | 0.000 | 0.000 |
| 1.014 | 0.000 | 0.000 | 1.011 | 0.000 | 0.000 | 1.008 | 0.000 | 0.000 | 1.011 | 0.000 | 0.000 | 1.010 | 0.002 | 0.001 |
| 1.026 | 0.000 | 0.001 | 1.022 | 0.002 | 0.003 | 1.017 | 0.003 | 0.001 | 1.019 | 0.001 | 0.001 | 1.018 | 0.003 | 0.001 |
| 1.041 | 0.001 | 0.000 | 1.033 | 0.004 | 0.006 | 1.025 | 0.006 | 0.006 | 1.030 | 0.003 | 0.006 | 1.029 | 0.005 | 0.005 |
| 1.056 | 0.004 | 0.003 | 1.044 | 0.006 | 0.012 | 1.032 | 0.007 | 0.011 | 1.041 | 0.006 | 0.013 | 1.039 | 0.005 | 0.010 |
| 1.068 | 0.004 | 0.005 | 1.055 | 0.010 | 0.017 | 1.042 | 0.015 | 0.019 | 1.052 | 0.007 | 0.019 | 1.049 | 0.007 | 0.018 |
| 1.083 | 0.012 | 0.007 | 1.066 | 0.018 | 0.025 | 1.049 | 0.023 | 0.037 | 1.062 | 0.010 | 0.028 | 1.058 | 0.013 | 0.030 |
| 1.099 | 0.021 | 0.015 | 1.078 | 0.024 | 0.038 | 1.057 | 0.038 | 0.054 | 1.073 | 0.019 | 0.047 | 1.068 | 0.019 | 0.049 |
| 1.112 | 0.026 | 0.019 | 1.089 | 0.039 | 0.058 | 1.067 | 0.055 | 0.088 | 1.084 | 0.024 | 0.076 | 1.079 | 0.023 | 0.075 |
| 1.127 | 0.048 | 0.030 | 1.101 | 0.057 | 0.088 | 1.075 | 0.077 | 0.125 | 1.096 | 0.038 | 0.118 | 1.090 | 0.034 | 0.110 |
| 1.144 | 0.074 | 0.044 | 1.113 | 0.088 | 0.116 | 1.083 | 0.107 | 0.169 | 1.105 | 0.049 | 0.156 | 1.099 | 0.048 | 0.150 |
| 1.157 | 0.103 | 0.058 | 1.124 | 0.121 | 0.154 | 1.093 | 0.148 | 0.234 | 1.117 | 0.069 | 0.221 | 1.109 | 0.057 | 0.206 |
| 1.174 | 0.155 | 0.080 | 1.137 | 0.160 | 0.195 | 1.101 | 0.186 | 0.300 | 1.129 | 0.091 | 0.297 | 1.121 | 0.073 | 0.283 |
| 1.190 | 0.222 | 0.102 | 1.149 | 0.209 | 0.244 | 1.109 | 0.235 | 0.382 | 1.141 | 0.118 | 0.394 | 1.132 | 0.089 | 0.373 |
| 1.204 | 0.300 | 0.131 | 1.161 | 0.264 | 0.298 | 1.119 | 0.298 | 0.493 | 1.151 | 0.147 | 0.493 | 1.141 | 0.111 | 0.471 |
| 1.221 | 0.413 | 0.167 | 1.174 | 0.335 | 0.359 | 1.127 | 0.365 | 0.604 | 1.163 | 0.185 | 0.635 | 1.152 | 0.135 | 0.609 |

## 5 Results

We train our Constitutive Artificial Neural Network from Figure 1 with the biaxial extension data from Tables 1 and 2, either individually for each dataset or simultaneously for all datasets combined. For each training case, in every direction, we compare the experimentally reported stress-stretch data to the discovered stress-stretch model and use the other cases for testing. We report the correlation coefficients *R*^2^ as an indicator for the goodness-of-fit for both train and test data.

### For insufficiently rich training data, model discovery is non-unique

Figure 2 shows the discovered models for the strip-x and strip-y data of rabbit skin in Table 1. The four columns show four different models for four *different initial conditions*; the two rows show the stress-stretch relations in the *x*– and *y*–directions. First and foremost, the neural network is able to discover models that explain the data with *R*^2^ values of the order of *R*^2^ = 0.85 and above, except for the last column. As a general trend, the network discovers free-energy functions with pairs of two terms: one term is a function of one of the isotropic invariants, *I*_1_ or *I*_2_ shown in the hot reddish colors, and the other is a function of one of the anisotropic invariants, *I*_4_ or *I*_5_, shown in the cold bluish colors. Interestingly, the model only discovers pairs of quadratic exponential terms, exp([*I*_1_ – 3]^2^) and exp([*I*_4_ – 3]^2^) in the first and fourth column, exp([*I*_1_ – 3]^2^) and exp([*I*_5_ – 3]^2^) in the second column, exp([*I*_2_ – 3]^2^) and exp([*I*_5_ – 3]^2^) in the third column, while all linear, quadratic, and linear exponential terms train to zero. We conclude that for training with a set of strip-x and strip-y data, the network is able to discover models that approximate the data well. However, model detection is ambiguous and sensitive to network initialization.

**Figure 2:**
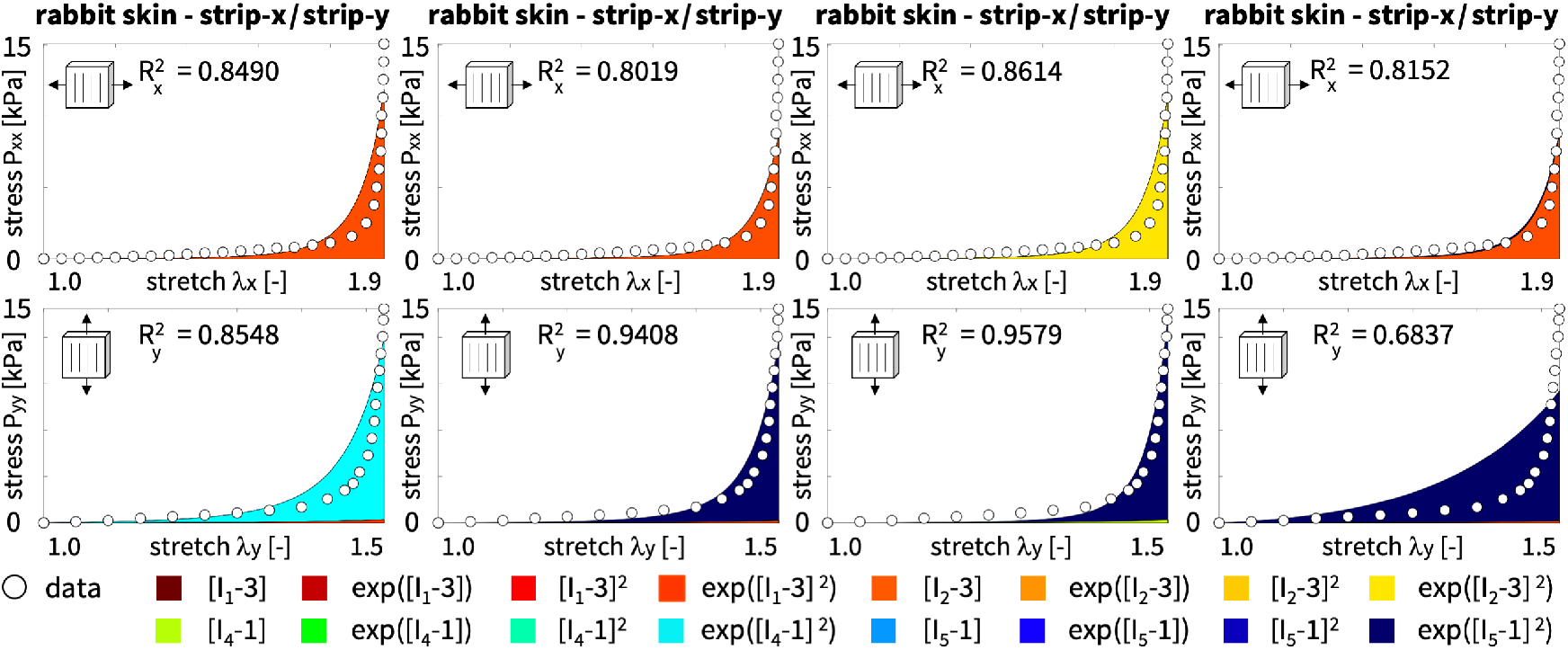
Rabbit skin strip-x and strip-y data and discovered model. Nominal stresses *P_xx_* and *P_yy_* as functions of stretches *λ_x_* at *λ_y_* = 1.000 and *λ_y_* at *λ_x_* = 1.000 for the transversely isotropic, perfectly incompressible Constitutive Artificial Neural Network with two hidden layers and 16 nodes from Figure 1, for training with *different initial conditions.* Dots illustrate the strip-x and strip-y data of rabbit skin [24] from Table 1; color-coded areas highlight the 16 contributions to the discovered stress function according to Figure 1.

This suggests that training with a single set of strip-x and strip-y data, with pairs of {*λ_x_*, *P_xx_*} and {*λ_y_*, *P_yy_*} from two independent experiments provides insufficient information for the discovery of unique models. To explore whether this ambiguity is inherent to the neural network itself or is simply due to insufficiently rich training data, we now train the model with multiple datasets from biaxial extension tests.

### For sufficiently rich training data, the network robustly discovers unique models

Figure 3 illustrates the discovered model for the biaxial extension data of rabbit skin in Table 1. The four columns show the discovered model for four different levels of lateral stretch, *λ_y_* = 1.000, 1.087, 1.235, 1.415, for training with each dataset *individually*; the two rows show the nominal stresses *P_xx_* and *P_yy_* for increasing stretches *λ_x_*. Similar to the previous example, the neural network is generally able to discover a model that explains the data reasonably well, with *R*^2^ values on the order of *R*^2^ =0.75 and above. Interestingly, for the four sets of biaxial extension data, even when trained on only one dataset, the neural network robustly discovers the same pair of terms: the quadratic exponential terms of the first invariant, exp([*I*_1_ – 3]^2^) in light red, and of the the fourth invariant, exp([*I*_4_ – 3]^2^) in turquoise. In the first column, the stretch perpendicular to the fiber direction, *λ_x_* = 2.0, is up to twice as large as the stretch parallel to the fiber direction, *λ_y_* = 1.0, and the behavior of the sample is dominated by the isotropic response of the light red exp([*I*_1_ – 3]^2^) term. With increasing fiber stretch, the anisotropic contribution of the fibers increases from the left to the right column. In the fourth column, the stretch perpendicular to the fiber direction, *λ_x_* = 1.5, is almost identical to the stretch parallel to the fiber direction, *λ_y_* = 1.4, and the behavior of the sample is dominated by the anisotropic response of the turquoise exp([*I*_4_ – 3]^2^) term. Interestingly, the second and fifth invariants *I*_2_ and *I*_5_ do not contribute to the discovered model, nor do the linear, quadratic, and linear exponential terms. From the robust activation of the same two terms across all datasets, we conclude that even a single individual set of biaxial training data with data triples of {*λ_x_*, *P_xx_*, *P_yy_*} allows for a more robust training than a set of strip-x and strip-y data, with pairs of {*λ_x_*, *P_xx_*} and {*λ_y_*, *P_yy_*} from two independent experiments. While the *parameter values are different* for each set of training data, the *set of active parameters is the same* across all four datasets and defines the discovered model. To check the robustness of the model findin, we now train our neural network simultaneously with all four biaxial extension datasets combined.

**Figure 3:**
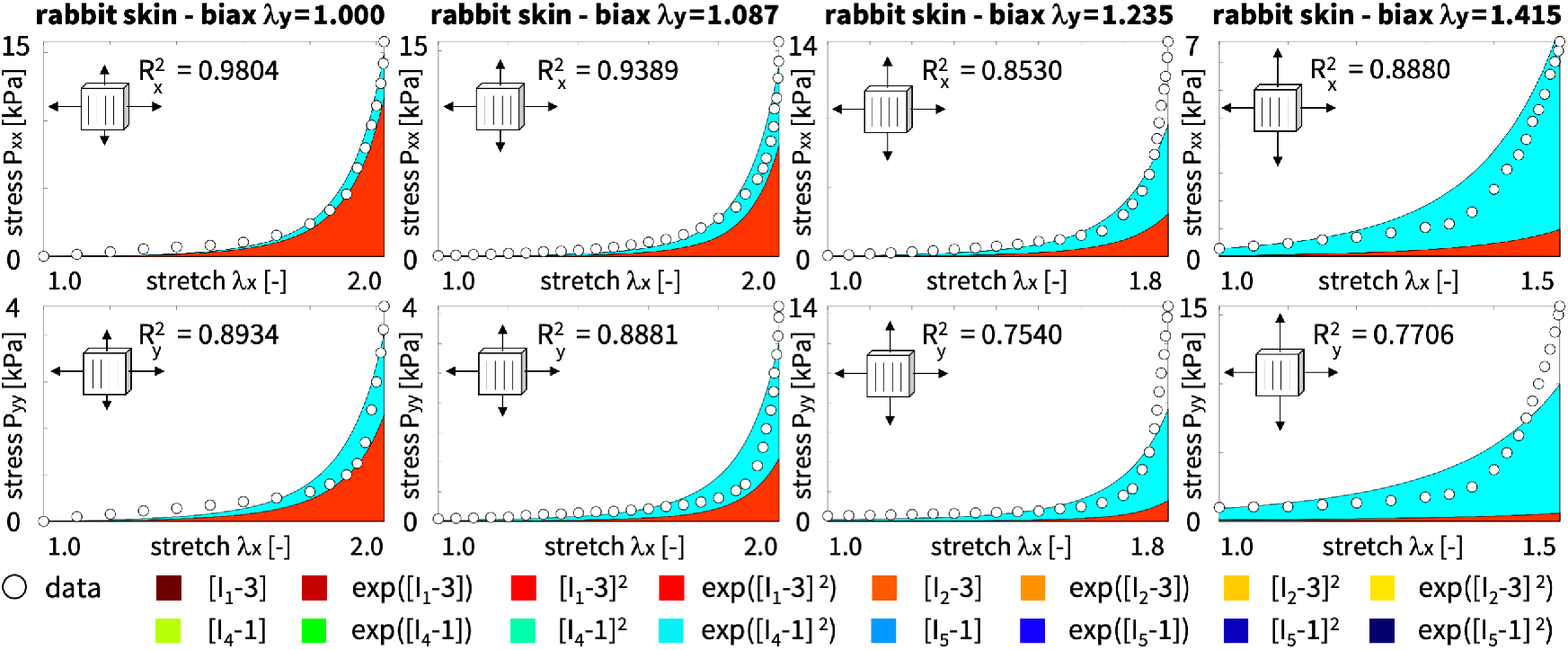
Rabbit skin biaxial extension data and discovered model. Nominal stresses *P_xx_* and *P_yy_* as functions of the lateral stretch *λ_x_* at *λ_y_* = 1.000, 1.087, 1.235, 1.415 for the transversely isotropic, perfectly incompressible Constitutive Artificial Neural Network with two hidden layers and 16 nodes from Figure 1, trained with each dataset *individually*. Dots illustrate the biaxial extension data of rabbit skin [24] from Table 1; color-coded areas highlight the 16 contributions to the discovered stress function according to Figure 1.

### The network autonomously discovers a two-term quadratic exponential model for rabbit skin

Figure 4 illustrates the discovered model for the biaxial extension data of rabbit skin in Table 1. The four columns show the response of the discovered model for four different levels of lateral stretch, *λ_y_* = 1.000, 1.087, 1.235, 1.415, but now for training with all four datasets *simultaneously;* the two rows show the nominal stresses *P_xx_* and *P_yy_* for increasing stretches *λ_x_*. When trained with four datasets of triples of {*λ_x_*, *P_xx_*, *P_yy_*}, the network robustly discovers a single unique model and parameter set. The model and parameters approximate the data well and are insensitive to varying initial conditions. The network consistently discovers a two-term model for rabbit skin in terms of quadratic exponentials of the first and fourth invariants *I*_1_ and *I*_4_, i.e.

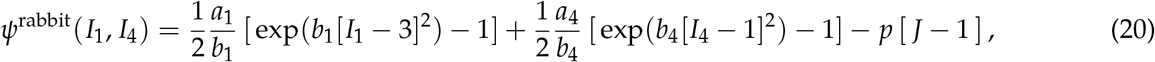

from which we can derive the nominal stress for rabbit skin, ***P*** = *∂ψ*/*∂**F***, as

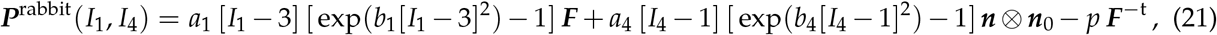

where ***n*** = ***F*** · ***n***_0_ and ***n***_0_ denote the deformed and undeformed collagen fiber directions, respectively. The network robustly discovers the same set of four non-zero network weights, *w*_1,4_, *w*_2,4_, *w*_1,12_, *w*_2,12_, that translate into four *physically interpretable parameters* with well-defined physical units, the stiffness-like parameters *a*_1_ = 2*w*_1,4_*w*_2,4_ and *a*_4_ = 2*w*_1,12_*w*_2,12_ and the unit-less exponential coefficients *b*_1_ = *w*_1,4_ and *b*_4_ =*w*_1,12_.

**Figure 4:**
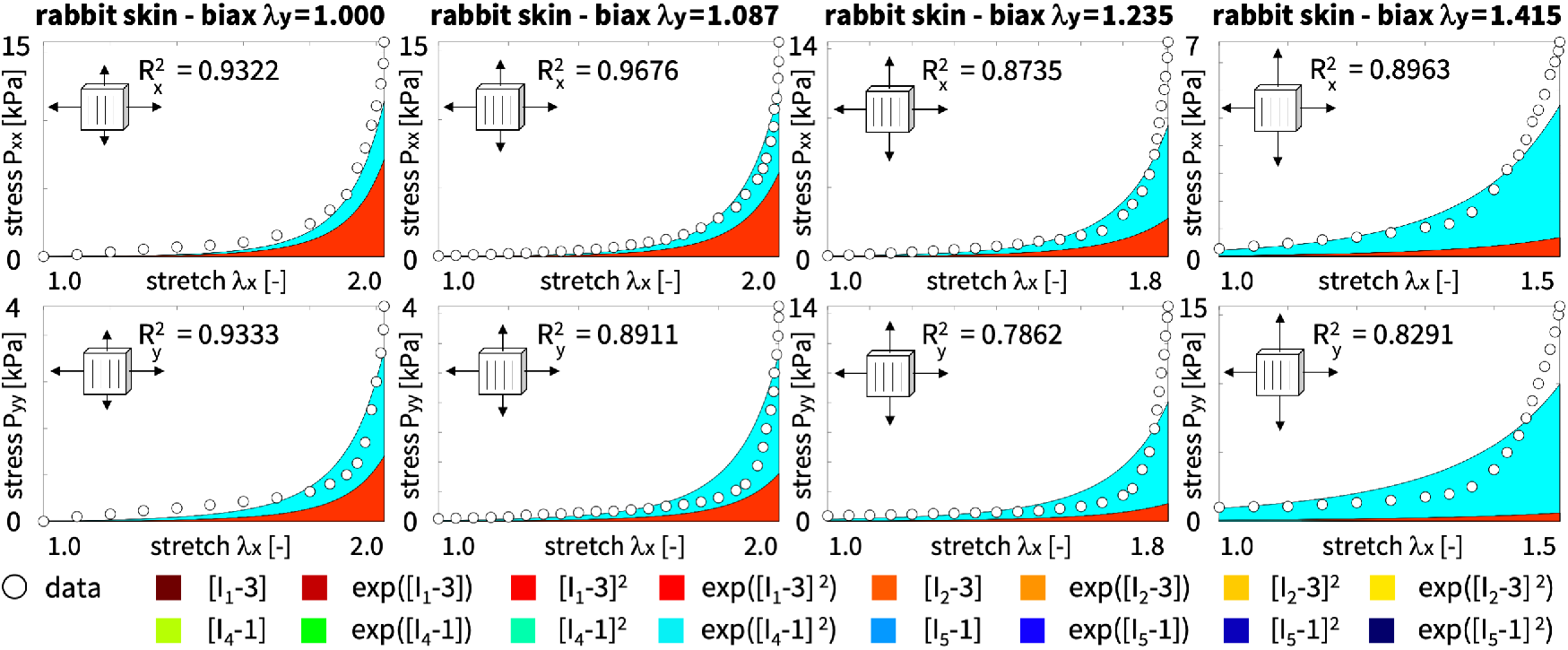
Rabbit skin biaxial extension data and discovered model. Nominal stresses *P_xx_* and *P_yy_* as functions of the stretch *λ_x_* at fixed *λ_y_* = 1.000, 1.087, 1.235, 1.415 for the transversely isotropic, perfectly incompressible Constitutive Artificial Neural Network with two hidden layers and 16 nodes from Figure 1, trained with all four datasets *simultaneously.* Dots illustrate the biaxial extension data of rabbit skin [24] from Table 1; color-coded areas highlight the 16 contributions to the discovered stress function according to Figure 1 for training with all four datasets combined.

Figure 5 summarizes the experimentally reported and computationally discovered stress-stretch relations for the biaxial extension of rabbit skin. The red curves, associated with the smallest fiber stretch of *λ_y_* = 1.000, are dominated by the isotropic response of the tissue and display the softest response. Increasing the fiber stretch from *λ_y_* = 1.000 via *λ_y_* = 1.087 and *λ_y_* = 1.235 to *λ_y_* = 1.415, from red to dark blue, gradually activates the anisotropic response of the collagen fibers and the stresses increase. The blue curves associated with the largest fiber stretch of *λ_y_* = 1.415 are dominated by the anisotropic response of the fibers and display the stiffest response in both directions. Overall, we conclude that the discovered two-term stress-stretch relation (21) provides a good approximation to the data for both stresses, in the stretch direction *P_xx_* and in the hold direction *P_yy_*. The model captures well the characteristic stretch stiffening of soft collagenous tissues with increasing stresses, as soon as the collagen fibers as the load-carrying structural element take over the main load. To validate the model discovery, we now train our neural network with a different dataset from biaxial extension experiments on pig skin.

**Figure 5:**
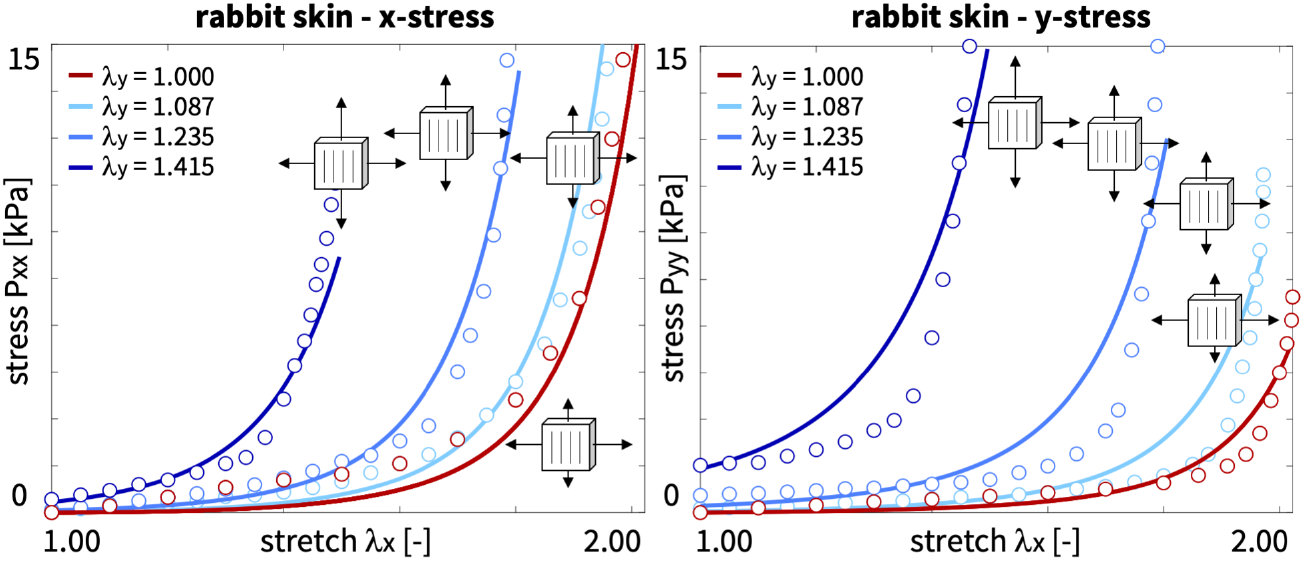
Rabbit skin biaxial extension tests and discovered model. Nominal stresses *P_xx_* and *P_yy_* as a function of stretch *λ_x_* at fixed *λ_y_* = 1.000, 1.087, 1.235, 1.415, from red to blue, for the transversely isotropic, perfectly incompressible Constitutive Artificial Neural Network with two hidden layers and 16 nodes from Figure 1, trained with all four datasets *simultaneously.* Dots illustrate the biaxial extension data of rabbit skin [24] from Table 1; colored curves highlight the discovered stress functions.

### The network discovers the same two-term model for rabbit and pig skin

Figure 6 illustrates the discovered model for the biaxial extension data of pig skin in Table 2. The four columns show the response of the discovered model for the strip-x and strip-y, off-x, equi-biaxial, and off-y tests; the two rows show the true stresses *σ_xx_* and *σ_yy_* as functions of the stretches *λ_x_* and *λ_y_*. When trained with the five stress-stretch pairs {*λ_x_*, *P_xx_*} and {*λ_y_*, *P_yy_*} *individually* the network robustly discovers the same model for each dataset. The model is insensitive to varying initial conditions and approximates the data well with *R*^2^ values on the order of *R*^2^ = 0.92 and above. Remarkably, even for a completely different dataset, from different species, tested with a different protocol, the network discovers the same model for pig skin and for rabbit skin half a century later with much higher precision: a two-term model in terms of quadratic exponential first and fourth invariants *I*_1_ and *I*_4_. The stress contributions to *σ_xx_* and *σ_yy_* in the two rows clearly visualize the effect of the collagen fibers: the *σ_xx_* stresses perpendicular to the fiber direction only contain the light red quadratic exponential isotropic *I*_1_ term, while the *σ_yy_* stresses parallel to the fiber direction also contain the turquoise quadratic exponential anisotropic *I*_4_ term. For all five datasets, only these two terms are activated, while the weights of the other 14 terms train to zero. While the *parameter values are different* for each set of training data, the *set of active parameters is the same* across all five datasets and defines the discovered model. Table 3 summarizes the non-zero weights *w*_1,4_, *w*_2,4_, *w*_1,12_, *w*_2,12_, the resulting stiffness-like parameters, *a*_1_ and *a*_4_, and exponential coefficients, *b*_1_ and *b*_4_, and the goodness-of-fit, 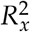 and 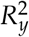, for individual training with the strip-x, off-x, equi-biaxial, off-y, and strip-y tests.

**Figure 6:**
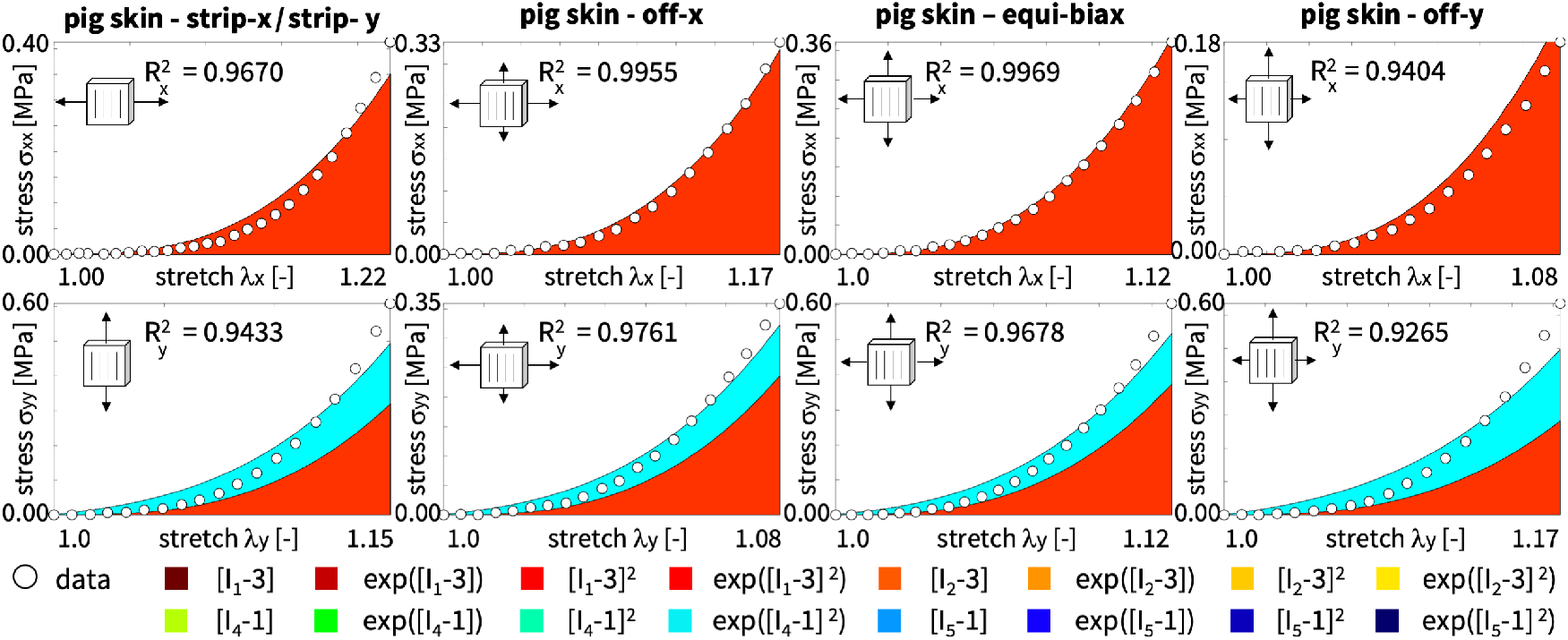
Pig skin biaxial extension data and discovered model. True stresses *σ_xx_* and *σ_yy_* as functions of stretches *λ_x_* and *λ_y_* for the transversely isotropic, perfectly incompressible Constitutive Artificial Neural Network with two hidden layers and 16 nodes from Figure 1, trained with each dataset *individually.* Dots illustrate the biaxial extension data of pig skin [51] from Table 2; color-coded areas highlight the 16 contributions to the discovered stress function according to Figure 1.

**Table 3:** Pig skin parameters from biaxial extension tests. Discovered material parameters for training with strip-x, off-x, equi-biaxial, off-y, strip-y tests from Table 2, trained with each dataset *individually,* parameter means, and trained with all five datasets *simultaneously.* Summary of the four non-zero weights *w*_1,4_, *w*_2,4_, *w*_1,12_, *w*_2,12_; resulting stiffness-like parameters *a*_1_ and *a*_4_ and exponential coefficients *b*_1_ and *b*_4_; and goodness-of-fit 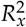 and 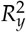.

|  | pig strip-x | pig off-x | pig equi-biax | pig off-y | pig strip-y | pig mean | pig all |
| --- | --- | --- | --- | --- | --- | --- | --- |
| | $\lambda_y = 1.00$ | $\lambda_y = \sqrt{\lambda_x}$ | $\lambda_y = \lambda_x$ | $\lambda_x = \sqrt{\lambda_y}$ | $\lambda_x = 1.00$ | | |
| $w_{1,4}$ [-] | 0.8085 | 0.7888 | 0.9400 | 0.8052 | 1.3075 | 0.9300 | 0.8207 |
| $w_{2,4}$ [MPa] | 0.7938 | 0.7770 | 0.9205 | 0.7872 | 1.3004 | 0.9158 | 0.8097 |
| $w_{1,12}$ [-] | 0.0006 | 0.3266 | 0.3332 | 0.3779 | 0.3429 | 0.2762 | 0.3921 |
| $w_{2,12}$ [MPa] | 0.0011 | 0.3226 | 0.3067 | 0.2832 | 0.2809 | 0.2389 | 0.3388 |
| $a_1$ [MPa] | 1.2837 | 1.2257 | 1.7304 | 1.2677 | 3.4004 | 1.7816 | 1.3291 |
| $b_1$ [-] | 0.8085 | 0.7888 | 0.9400 | 0.8052 | 1.3075 | 0.9300 | 0.8207 |
| $a_4$ [MPa] | 0.0000 | 0.2107 | 0.2044 | 0.2140 | 0.1927 | 0.1644 | 0.2656 |
| $b_4$ [-] | 0.0006 | 0.3266 | 0.3332 | 0.3779 | 0.3429 | 0.2762 | 0.3921 |
| $R_x^2$ [-] | 0.9945 | 0.9404 | 0.9969 | 0.9955 | 0.9670 | — | — |
| $R_y^2$ [-] | 0.9433 | 0.9265 | 0.9678 | 0.9761 | 0.9764 | — | — |

### The network simultaneously discovers both a unique model and parameter set

Figure 7 confirms the robust model discovery, now for training with all five datasets *simultaneously*. The network discovers the same two-term model for pig skin as in the previous example for rabbit skin, with the same free-energy function as in eq. (20), i.e.

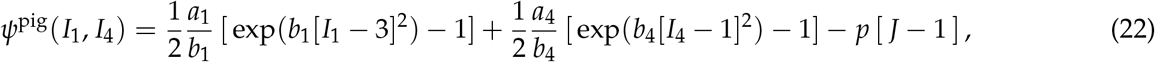

from which we can derive the true stress, ***σ*** = 1/*J*∂*ψ*/∂***F*** · ***F*^t^** = 1/*J* ***P*** · ***F***^t^ similar to eq. (21), i.e.

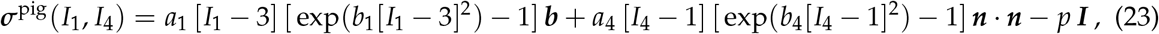

where ***n*** = ***F*** · ***n***_0_ denotes the deformed collagen fiber direction. Simultaneous training with all five combined datasets confirms that the second and fifth invariants *I*_2_ and *I*_5_ do not contribute to the discovered model, nor do the linear, quadratic, and linear exponential terms. In contrast to the individual training, the simultaneous training results in a *single unique set of parameter values* that best explain all five experiments combined. Naturally, simultaneous training slightly reduces the goodness-of-fit *R*^2^ compared to individual training. Importantly, the four non-zero weights of the discovered model, *w*_1,4_ = 0.8207, *w*_2,4_ = 0.8097 MPa, *w*_1,12_ = 0.3921, *w*_2,12_ = 0.3388 MPa, naturally translate into as set of meaningful, *physically interpretable parameters*, *a*_1_ = 2*w*_1,4_*w*_1,4_ = 1.3291MPa, *b*_1_ = *w*_1,4_ = 0.8207, *a*_4_ = 2*w*_1,12_*w*_1,12_ = 0.2656MPa, and *b*_4_ = *w*_1,12_ = 0.3921 with real physical units. Table 3 compares the discovered non-zero weights *w*_1,4_, *w*_2,4_, *w*_1,12_, *w*_2,12_, the resulting stiffness-like parameters *a*_1_ and *a*_4_ and exponential coefficients *b*_1_ and *b*_4_, and goodness-of-fit 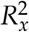 and 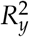 for simultaneous training to the means of their counterparts for individual training. Figure 8 summarizes the experimentally reported and computationally discovered stress-stretch relations for biaxial extension of pig skin. The red curves of the strip-x test with *λ_y_* = 1.000 are dominated by the isotropic response of the tissue and display the softest response. Increasing the fiber stretch from *λ_y_* =1.000 via 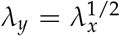, *λ_y_* = *λ_x_*, and 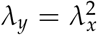, to the strip-y test with *λ_x_* = 1.000, from red to dark blue, gradually activates the anisotropic response of the collagen fibers and the stresses increase. The dark blue curves are dominated by the anisotropic response of the fibers and display the stiffest response. Similar to rabbit skin, the discovered two-term stress-stretch relation for pig skin (23) provides a good approximation of the data for the stresses in both *σ_xx_* and *σ_yy_* directions. We conclude that the discovered model is not only well-suitable for rabbit skin, but also for pig skin.

**Figure 7:**
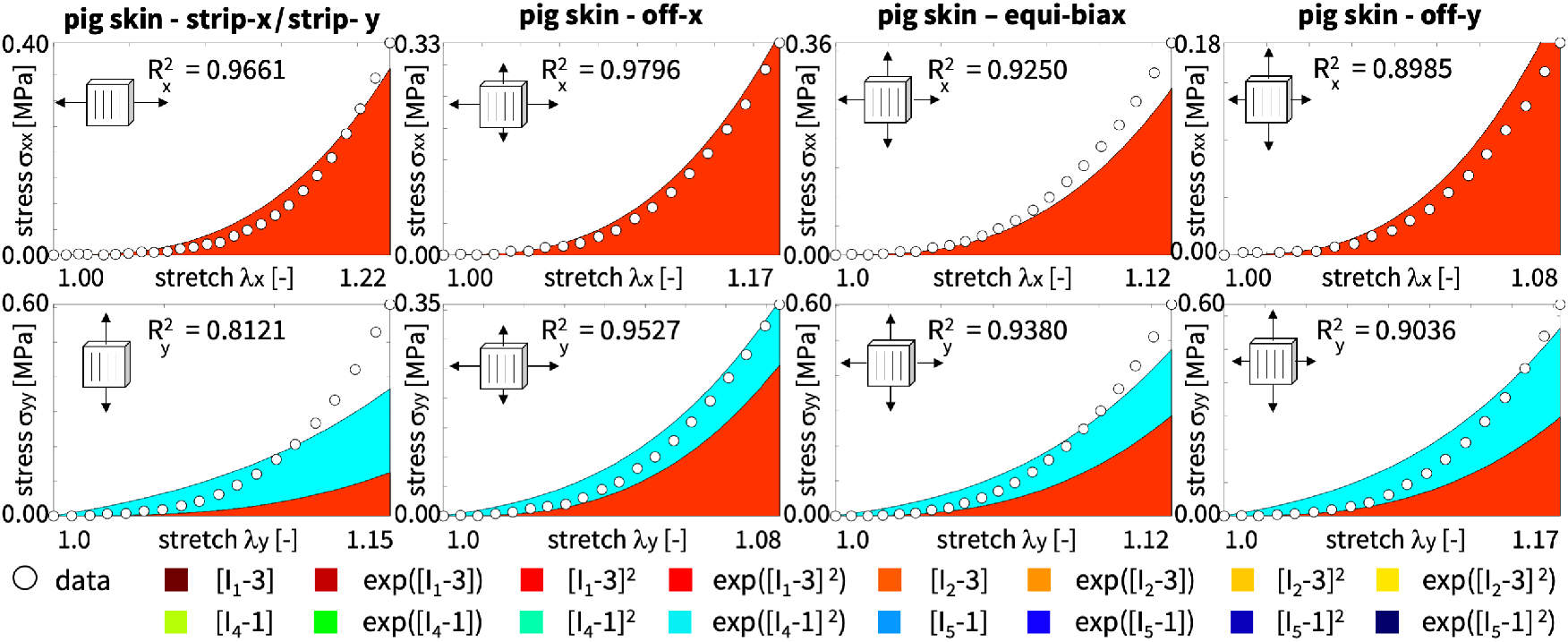
Pig skin biaxial extension data and discovered model. True stresses *σ_xx_* and *σ_yy_* as functions of the stretches *λ_x_* and *λ_y_* for the transversely isotropic, perfectly incompressible Constitutive Artificial Neural Network with two hidden layers and 16 nodes from Figure 1, trained with all five datasets *simultaneously*. Dots illustrate the biaxial extension data of pig skin [51] from Table 2; color-coded areas highlight the 16 contributions to the discovered stress function according to Figure 1.

**Figure 8:**
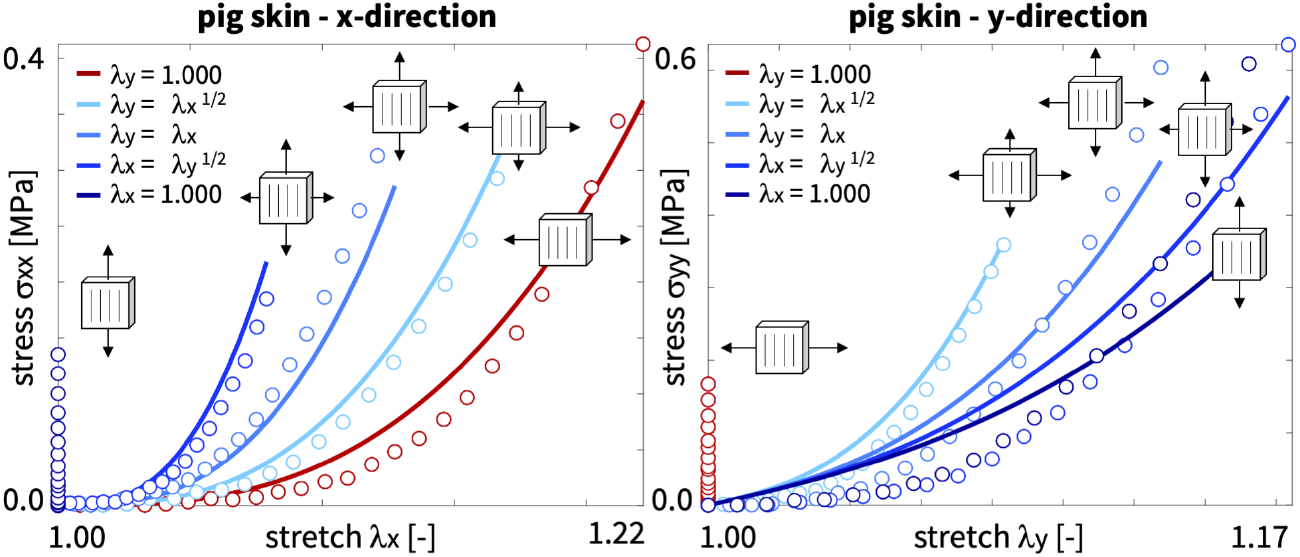
Pig skin biaxial extension tests and discovered model. True stresses *σ_xx_* and *σ_yy_* as functions of stretches *λ_x_* and *λ_y_* for the transversely isotropic, perfectly incompressible Constitutive Artificial Neural Network with two hidden layers and 16 nodes from Figure 1, trained with all five dataset *simultaneously.* Dots illustrate the biaxial extension strip-x, off-x, equibiaxial, off-y, strip-y data of pig skin [51] from Table 2; colored curves highlight the discovered stress functions.

### The network not only discovers the best model and parameters, but also the best experiment

Figure 9 visualizes the coefficients of correlation for the pig skin experiments. The six columns correspond to the six sets of training data, strip-x, off-x, equibiaxial, off-y, trip-y, and all experiments combined; the two rows illustrate the fit in the *x*– and *y*–directions. The color-coded bars summarize the *R*^2^ values from each experiment; the highlighted bar indicates the goodness-of-fit to the training data, 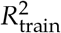, while all other bars indicate the goodness-of-fit to the test data, 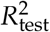. The discovered model clearly trains well, with all but two values well above 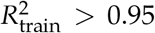. As expected, the strip-x experiment has no predictive potential for the *y*–stresses *σ_yy_* and neither does the strip-y experiment for the *x*–stresses *σ_xx_*, both with 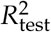 values of zero. At the same time, the off-x, equi-biaxial, and off-y experiments contain information about both stresses, *σ_xx_* and *σ_yy_*, and provide insight into both, the isotropic *I*_1_ term and the anisotropic *I*_4_ term, where most of the 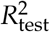 values are close to 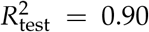. Importantly, comparing the mean coefficient of correlation, mean(*R*^2^), across all five experiments helps us select the experiment with the richest information: the mean coefficient of correlation is largest for the off-y and off-x experiments with values of mean(*R*^2^) = 0.8510 and 0.8423 and smallest for the strip-y experiment with mean(*R*^2^) = 0.2519. We conclude that our network can not only discover the best model and parameters, but also *discover the best experiment.* In other words, if we could choose just one experiment, the off-y experiment with 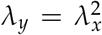 would provide the most complete information about the mechanics of skin.

**Figure 9:**
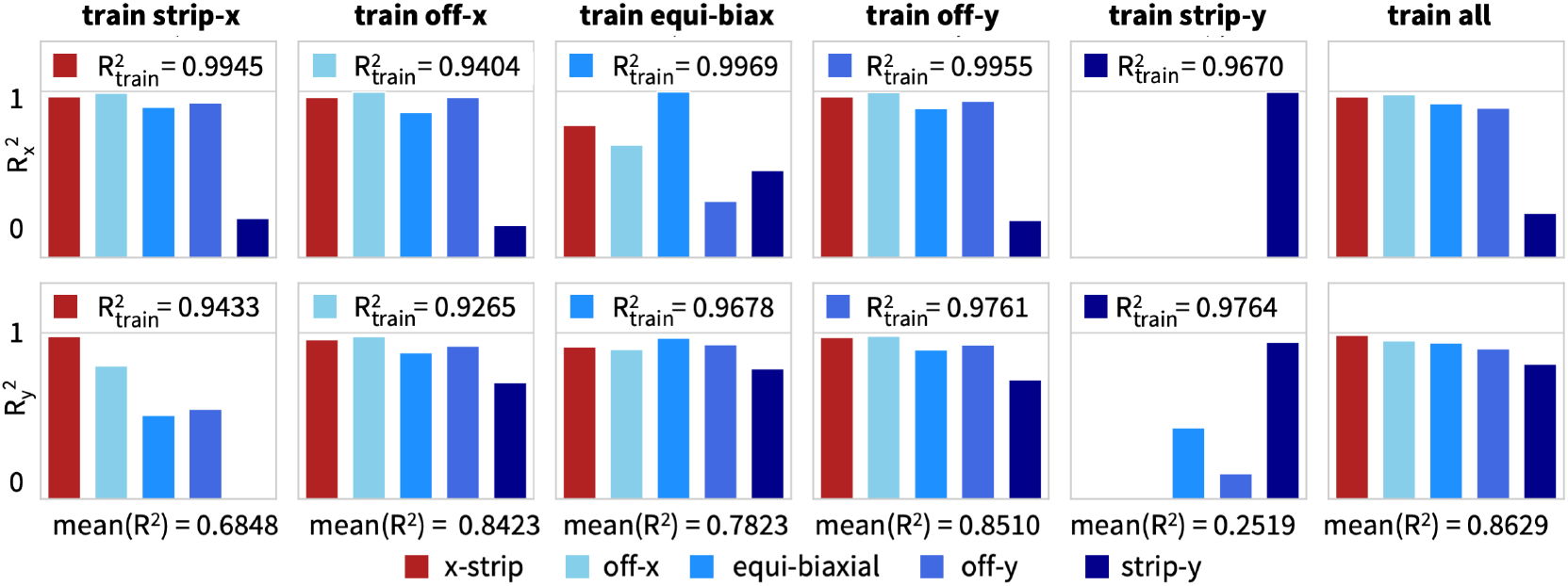
Pig skin biaxial extension coefficients of correlation. Coefficients of determination *R*^2^ for *discovered model,* with quadratic exponential terms in the first and fourth invariants, exp([*I*_1_ – 3]^2^) and exp([*I*_4_ – 1]^2^), trained with the strip-x, off-x, equibiaxial, off-y, trip-y, and all data of pig skin [51] from Table 2. Color-coded bars highlight the individual experiment. In each column, one experiment is used as training data and the other four as test data. In the right column, all data are used as training data.

### The network can discover microstructural features

Skin is a transversely isotropic material with a pronounced collagen fiber direction. If we do not know the fiber direction a priori, we can discover it simultaneously with the model and its parameters. We assume that the fibers are oriented at an angle *α* to the *x*-direction such that ***n***_0_ = [cos(*α*),sin(ff),0]^t^. Figure 1 of our Constitutive Artificial Neural Network introduces the fiber angle *α* as an additional weight *w*_0_ between the deformation gradient ***F*** and the fourth and fifth invariants, *I*_4_ = ***F*** · ***n***_0_ ⊗ ***n***_0_ · ***F***^t^ and *I*_5_ = ***F***^t^ · ***F*** · ***n***_0_ ⊗ ***n***_0_ · ***F***^t^ · ***F***. For the rabbit skin experiments in Table 1, the model discovers a weight of *w*_0_ = 0.9760 corresponding to an angle of *α* = ±55.92^0^; for the pig skin experiments in Table 2 the model discovers a weight of *w*_0_ = 1.5708 corresponding to an angle of *α* = 90.00^0^. Both values seem reasonable for skin tissue samples taken from the back of the animals with a pronounced lateral fiber orientation. Intuitively, the rabbit is probably too small to provide 30 × 30 mm^2^ tissue samples with homogeneous collagen fiber orientations that could explain why the learned fiber angle deviates from the lateral direction. However, the pig is much larger and its samples might be more homogeneous with a single unique collagen fiber orientation along the 90^0^ lateral direction. Overall, we conclude that, given sufficient training data, the network robustly *discovers microstructural features,* e.g., distinct collagen fiber directions of soft biological tissues.

## 6 Discussion

The objective of this study was to design a Constitutive Artificial Neural Network for transversely isotropic perfectly incompressible materials and to demonstrate its features using the example of skin. Our design paradigm was to reverse-engineer the network architecture to ensure that the network satisfies common physical and thermodynamic constraints by design and includes popular constitutive models as special cases. We have shown that a basic set of 16 functional building blocks–generated from two isotropic and two anisotropic invariants, their first and second powers, and their exponentials-provides a reasonable basis to characterize this class of materials. To leverage the robustness and stability of optimization tools in neural network modeling, we have integrated these terms into a neural network with two hidden layers and 32 weights. When the network is trained with biaxial extension data from the skin, it autonomously discovers a subset of non-zero weights that define the discovered model while training the majority of the weights to zero. Importantly, in contrast to classical neural network modeling, the non-zero weights are physically interpretable and translate naturally into engineering parameters and microstructural features such as shear modulus and fiber angle. The method not only discovers a unique model and parameter set that best describe the data, but also autonomously discovers the richest experiment to train itself.

### Our discovered model compares well against proposed constitutive models for skin

For more than half a century scientists have developed constitutive models to characterize the stress-stretch relation in skin [10, 19, 26, 30, 52, 59]. We can classify these models into microstructurally-based and invariant-based approaches [29]. Our neural network in Figure 1 with the free-energy function in eq. (10) is a natural generalization of the most popular invariant-based models [33] and provides insight into their functional correlations:

The *Lanir model* [27], the simplest of all models for transversely isotropic materials, has a free-energy function that contains an isotropic linear neo-Hookean term [55] of the first invariant [*I*_1_ – 3] and an anisotropic linear term of the fourth invariant [*I*_4_ – 1], i.e.

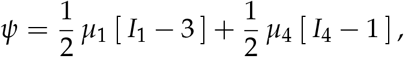

scaled by the shear modulus *μ*_1_ = 2*w*_1,1_*w*_2,1_ and the fiber stiffness *μ*_4_ = 2*w*_1,9_*w*_2,9_. While the Lanir model captures well the transversely isotropic behavior of skin with stiff collagen fibers embedded in a soft matrix, this is evident from the experimental stress-stretch curves in Figures 5 and 8. The linear neo-Hookean format does not capture the characteristic stretch-stiffening behavior of collagenous tissues [30].

The *Weiss model* [60] originally designed for ligaments combines the isotropic linear Mooney-Rivlin terms [40, 45] of the first and second invariant [*I*_1_ – 3] and [*I*_2_ – 3] with an anisotropic linear exponential Demiray term [11] of the fourth invariant [*I*_4_ – 1], i.e.

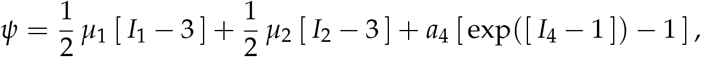

scaled by the shear moduli, *μ*_1_ = 2*w*_1,1_*w*_2,1_ and *μ*_2_ = 2*w*_1,5_*w*_2,5_, and the stiffness-like parameter *a*_4_ = 2*w*_1,10_*w*_2,10_. While the exponential format of the anisotropic term naturally captures the stretch-stiffening parallel to the collagen fiber direction, the linear format of the isotropic term does not capture the stretch-stiffening of the red curves perpendicular to the fiber direction in Figures 5 and 8.

The *Groves model* [15], a generalization of the Weiss model, combines an isotropic exponential term in the first invariant [*I*_1_ – 3] and an isotropic linear Blatz and Ko term [6] of the second invariant [*I*_2_ – 3] with an anisotropic linear exponential Demiray term [11] of the fourth invariant [*I*_4_ – 1], i.e.

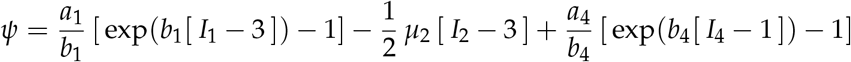

scaled by the stiffness-like parameters, *a*_1_ = 2*w*_1,2_*w*_2,2_, *μ*_2_ = 2*w*_1,5_*w*_2,5_, and *a*_4_ = 2*w*_1,10_*w*_2,10_, and the coefficients *b*_1_ = *w*_1,2_ and *b*_4_ = *w*_1,10_. While the exponential format of both the isotropic and anisotropic terms qualitatively captures the characteristic stretch-stiffening of the skin, the linear dependence in the exponential term accounts for only moderate stiffening and does not quantitatively capture the characteristic J-shape, particularly of the blue curves in Figures 5 and 8.

The *Holzapfel model* [17], combines the isotropic linear neo-Hookean term [55] of the first invariant [*I*_1_ – 3] with an anisotropic quadratic exponential term of the fourth invariant [*I*_4_ – 1], i.e.

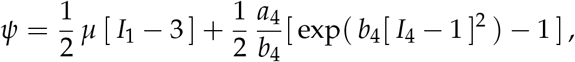

scaled by the shear modulus *μ* = 2*w*_1,1_*w*_2,1_, the stiffness-like parameter *a*_4_ = 2*w*_1,12_*w*_2,12_, and the coefficient *b*_4_ = *w*_1,12_. With only three parameters and a clear microstructural interpretation, the Holzapfel model naturally captures the three characteristic features of collagenous tissues, anisotropy, stretch-stiffening and a strong J-type behavior, and is probably the most popular model for soft biological tissues to date [18].

### Our network discovers interpretable parameter values that agree well with the values for skin

Instead of following the usual paradigm to *first* select a constitutive model and *then* identify its material parameters [29], our network discovers *simultaneously* both model and parameters. Notably, we offer the network a wide variety of functional building blocks [33] from which a relevant subset of two can be selected, i.e.

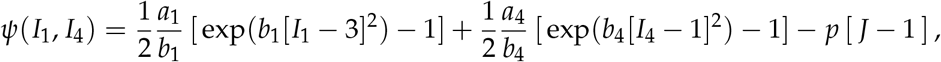

while all other network parameters train to zero. Unlike in classical neural networks, for which the weights have no physical interpretation [50, 51], our non-zero weights *w*_0_ = 1.5708, *w*_1,4_ = 0.8207, *w*_2,4_ = 0.8097 MPa, *w*_1,12_ = 0.3921, *w*_2,12_ = 0.3388 MPa, naturally translate into a set of meaningful, physically interpretable parameters: a collagen fiber angle of *α* = 90^0^, matrix and fiber stiffnesses of *a*_1_ = 1.3291 MPa and *a*_4_ = 0.2656 MPa, and matrix and fiber coefficients of *b*_1_ = 0.8207 and *b*_4_ = 0.3921, that can teach us something about the underlying microstructure and physics of skin [30].

### A special application of our neural network is parameter identification

By constraining the majority of weights to zero and only training for a selective subset of weights [34], we can utilize our neural network to identify the parameters of common constitutive models, including the Lanir [27], Weiss [60], Groves [15], or Holzapfel [17] models. For example, by training specifically for the weights *w*_1,1_, *w*_2,1_, *w*_1,12_, *w*_2,12_, we recover the classical Holzapfel parameters. For a simultaneous training with all five load cases from Table 2, the network discovers weights of *w*_1,1_ = 0.3240, *w*_2,1_ = 0.3845 MPa, *w*_1,12_ = 10.7914, and *w*_2,12_ = 0.0049 MPa that translate into a shear modulus of *μ* = 0.2492 MPa, a Holzapfel stiffness-like parameter of *a*_4_ = 0.1054 MPa, and a coefficient of *b*_4_ = 10.7914. Figures 10 and 11 summarize the training of our neural network when restricted to the classical Holzapfel model, [17] and trained with the biaxial extension data from pig skin in Table 2. Consistent with our intuition, the stress plots in Figure 10 confirm that the Holzapfel model performs well in the *y*–direction parallel to the primary collagen fiber orientation, but does not capture strain stiffening in the *x*-direction, perpendicular to the fibers. The coefficients of correlation in Figure 11 indicate an excellent fit for the off-y and strip-y data in the *y*-direction, but a moderate fit for all other data. Compared to the popular and widely used Holzapfel model with linear and quadratic exponential terms in Figures 10 and 11, our discovered model with two quadratic exponential terms in Figures 7 and 9 has larger overall coefficients of correlation *R*^2^ and provides a better fit of the data.

**Figure 10:**
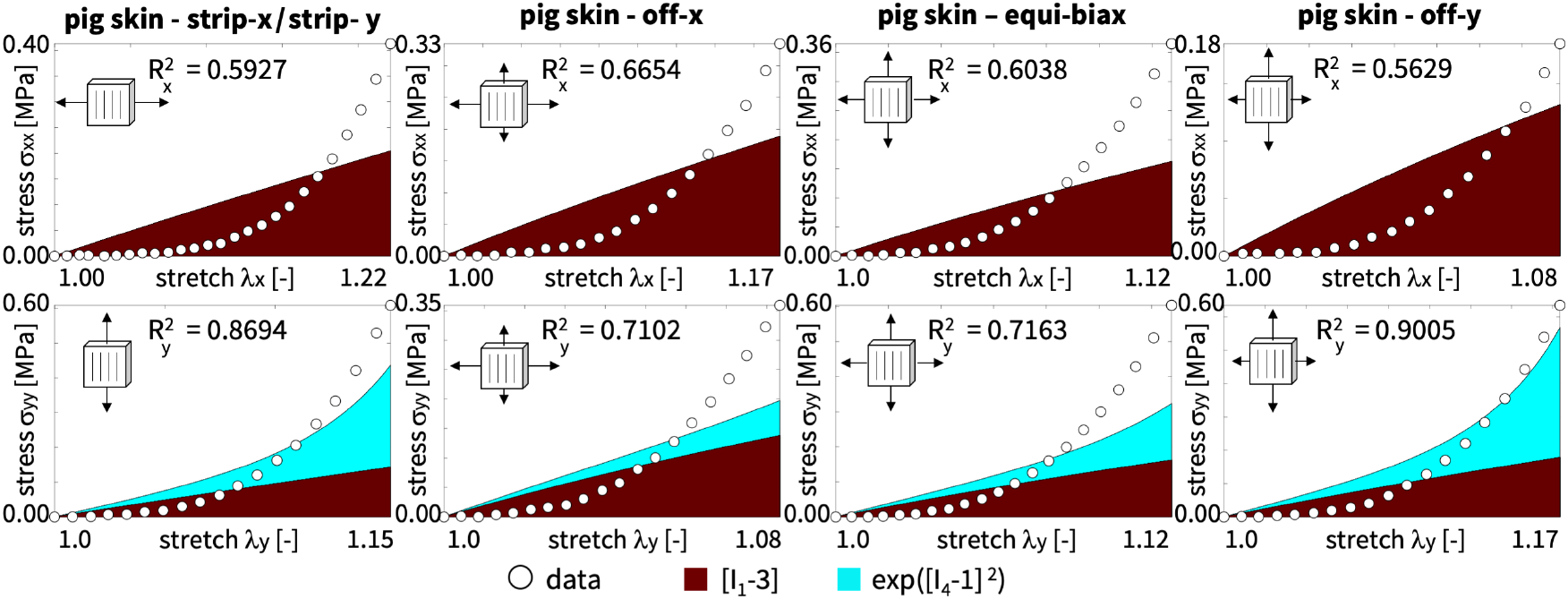
Pig skin biaxial extension data and neo-Hooke-Holzapfel model. True stresses *σ_xx_* and *σ_yy_* as functions of the stretches *λ_x_* and *λ_y_* for the *neo-Hooke-Holzapfel model,* trained with all five dataset *simultaneously.* Dots illustrate the biaxial extension data of pig skin [51] from Table 2; color-coded areas highlight the neo-Hookean and the Holzapfel contributions to the stress function.

**Figure 11:**
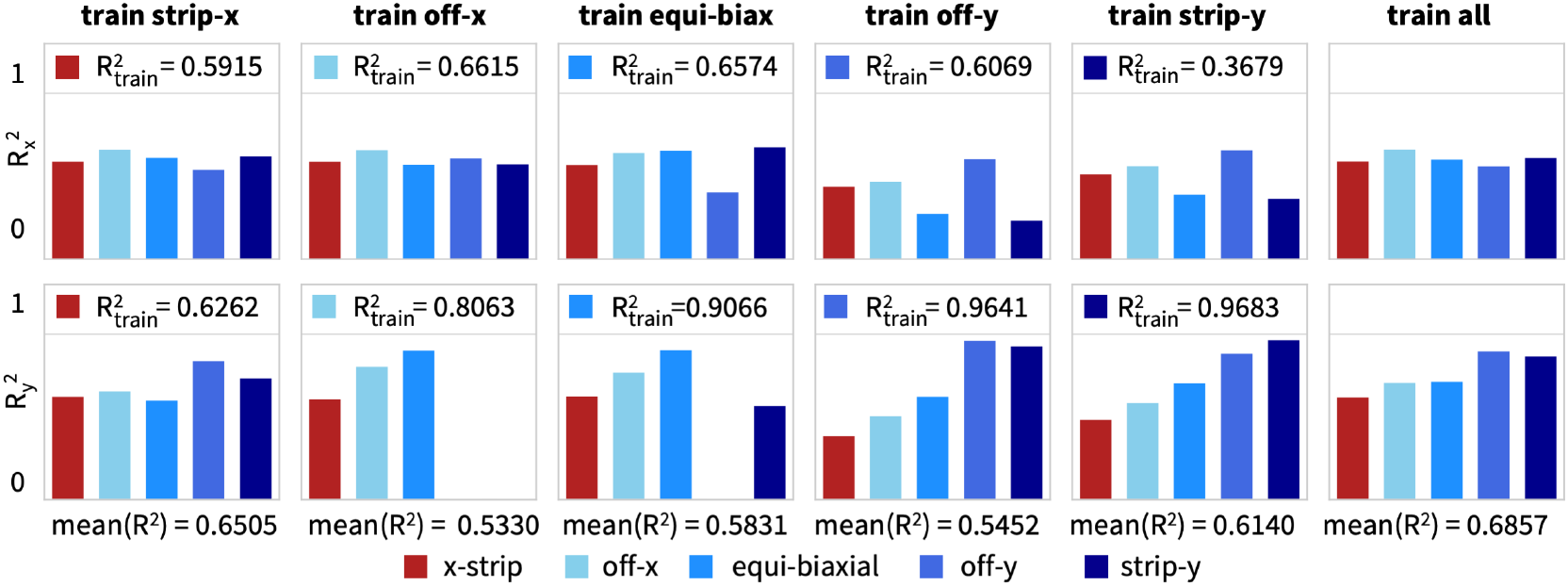
Pig skin biaxial extension coefficients of correlation. Coefficients of determination *R*^2^ for *neo-Hooke-Holzapfel model,* with linear and quadratic exponential terms in the first and fourth invariants, [*I*_1_ – 3] and exp([*I*_4_ – 1]^2^), trained with strip-x, off-x, equibiaxial, off-y, strip-y, and all data of pig skin [51] from Table 2. Color-coded bars highlight each experiment. In each column, one experiment is used as training data and the other four as test data. In the right column, all data are used as training data.

To address the shortcomings in the isotropic response of the classical Holzapfel model [17] in Figures 10 and 11, the *new Holzapfel model* [13] accounts for a fiber dispersion around the pronounced direction **n**_0_ with an additional dispersion parameter *κ*. It includes both the first and fourth invariants I1 and I4 in the quadratic exponential while preserving the isotropic linear neo-Hookean term, i.e.

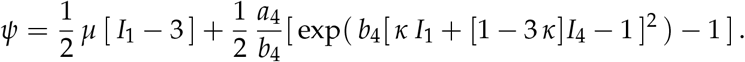

While our current neural network architecture is not fully connected and–by design–contains *explicit coupling between individual invariants*, it discovers two exponential quadratic terms that are formally similar to the exponential term in the new Holzapfel model. In contrast to the Holzapfel model, the *w*_1,1_ and *w*_2,1_ weights of the isotropic linear neo-Hookean term, 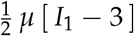, consistently train to zero. This indicates that in our parameterization the isotropic linear term plays a negligible role and all isotropic contributions can be collectively represented by our exponential quadratic term, 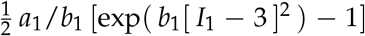, through non-zero *w*_1,4_ and *w*_2,4_ weights alone. Importantly, for a dispersion parameter of 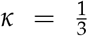, the new Holzapfel model [13] exactly recovers our exponential quadratic isotropic term, while for *κ* = 0, it recovers our exponential quadratic anisotropic term, 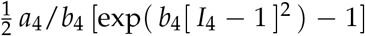. These direct comparisons with classical constitutive models [13, 15, 17, 27, 60] suggests that our network not only discovers a physically reasonable model with a small number of well-rationalized functional building blocks, but also helps rationalize the features and shortcomings of existing models with regard to the development history of constitutive models [19, 29, 30].

### Our neural network is polyconvex by design

From the general representation theorem [46] we know that the free-energy function of an isotropic material can be expressed in its most generic form as an infinite series of power products of its invariants, i.e. 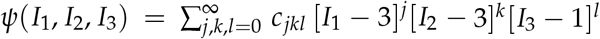, where *c_jkl_* are material constants. It is easy to see that the format of this function is more general than our free-energy function (10): it contains *mixed products of invariants*, for which the free-energy function is generally not convex. However, we can design the free-energy function as a *sum of convex functions of invariants* such that the overall free-energy function remains polyconvex [16]. This has motivated us to represent the free energy as the sum of four individual polyconvex subfunctions *ψ*_1_, *ψ*_2_, *ψ*_4_, *ψ*_5_ [2, 33], such that *ψ*(***F***) = *ψ*_1_ (*I*_1_) + *ψ*_2_(*I*_2_) + *ψ*_4_(*I*_4_) + *ψ*_5_(*I*_5_), is polyconvexby design. For our neural network, this implies that instead of using a *fully connected network* architecture, in which coupling terms in the invariants *I*_1_, *I*_2_, *I*_4_, *I*_5_ emerge naturally, we propose a *selectively connected network* architecture in which the four inputs *I*_1_, *I*_2_, *I*_4_, *I*_5_ remain decoupled at all times, and combine only additively to the final free-energy function, *ψ* = *ψ*_1_ + *ψ*_2_ + *ψ*_4_ + *ψ_5_*, after the last hidden layer [21, 51]. In other words, if we want to include an explicit coupling of the invariants, e.g., through the dispersion term [*κI*_1_ + [1 – 3*κ*] *I*_4_ – 1] of the new Holzapfel model [13], we would have to extend our current network by adding connections between the first and second layers, but then we might have to add alternative strategies to ensure polyconvexity [12].

### Our network autonomously discovers the best experiment with the largest correlation coefficient

For almost half a century, scientists have proposed different experiments to characterize flat collagenous tissues, including uniaxial tension [41], biaxial extension [24, 39], torsion [5], suction [58], bulging [53], and indentation [19, 59]. By far the simplest method is plain uniaxial tension. Intuitively we would think that testing the sample *parallel* and *perpendicular* to the collagen fiber orientation, e.g., at angles of 0^0^,45^0^,90^0^, should provide the best insight into its overall response [41]. However, as we have seen in Figure 2, from single {*λ_x_*, *P_xx_*} and {*λ*_y_, *P_yy_*} stretch-stress pairs alone, it is difficult to interpret the complex anisotropic behavior of skin. The first pioneering biaxial test system for skin was proposed almost five decades ago [23], and has since then become the method of choice to characterize flat composite materials with stiff fibers embedded in a soft matrix. Instead of data pairs, this system provides data triplets, {*λ_x_*, *σ_xx_*, *σ_yy_*} and {*λ*_y_, *σ_xx_*, *σ_yy_*}, where the second stretch *λ_y_* or *λ_x_* is either held constant or increased as a function of *λ_x_* or *λ_y_* [24, 25, 50, 51]. From Figure 3 for rabbit skin and Figure 6 for pig skin, we conclude that this method provides rich enough data to discover both a unique model and a parameter set, even from single experiments. Interestingly, analytical optimization of biaxial test protocols reveals that two uniaxial stretch tests at mutually normal directions at 22.5^0^, 67.5^0^ provide richer information than tests performed at 0^0^,45^0^,90^0^ [28]. This agrees well with our correlation coefficients in Figure 9, for which the *off-x and off-y experiments* at 22.5^0^, 67.5^0^ are the best correlations across all five experiments with mean(*R*^2^) values of 0.8423 and 0.8510. This suggests that our method is able to autonomously discover the experiment that provides the best information to train itself.

### Our Constitutive Artificial Neural Networks interpolate, extrapolate, and explain constitutive behavior

Unlike traditional neural networks, which do not require any prior physics knowledge to interpolate data within a well-defined window [1], Constitutive Artificial Neural Networks explicitly modify the network input, output, and architecture to incorporate physical constraints into the network design [31]. This allows them to interpolate and extrapolate the stress-stretch response *within and beyond* a known stretch regime. Interestingly, the first neural network to approximates incremental principal strains in concrete from given stress increments, stresses, and strains is more than three decades old [14]. In the early days, neural networks served primarily as black box regression operators without accounting for physical considerations or thermodynamic constraints. Now there is a strong push in constitutive modeling to ensure that neural networks satisfy these constraints a priori [2, 21, 31, 37, 51]. The first family of Constitutive Artificial Neural Networks designed with these goals uses multiple hidden layers to map the deformation gradient to a free-energy function, from which they derive the stress [31]. Their layers are densely connected by conventional activation functions, typically of hyperbolic tangent or logistic type [33]. This introduces hundreds if not thousands of network weights and biases, which the network has to learn from data. Of course, with so many degrees of freedom, these early Constitutive Artificial Neural Networks have excellent interpolation properties: they can fit any stress-stretch curve flawlessly, including potential measurement outliers [51]. However, with more unknowns than data, they have a clear tendency of overfit [1]. Importantly, simply measuring more points along the same stress-stretch curve or testing more samples with the same protocol does not fix the overfitting problem. For successful training, the network does not simply need *more data*, but *rich data*, ideally from multi-mode tests, not only individually in uniaxial tension, biaxial extension, and torsion, but also at different fiber angles [50], and combined with all modes [7, 34]. Even with the best of all training data, two essential limitations remain: the lack of extrapolation and the lack of interpretability. Many popular conventional activation functions, such as the hyperbolic tangent or logistic functions, tend to plateau beyond a certain range, and this plateau naturally translates into the approximated stress function [33]. Furthermore, their weights are typically non-unique, lack clear physical interpretation, and offer only limited insight into the underlying constitutive response. Our new family of Constitutive Artificial Neural Networks addresses these limitations through a simple design paradigm: it reverse-engineers its activation functions from functional building blocks of well-accepted and widely used constitutive models, including the isotropic neo-Hookean [55], Blatz and Ko [6], Mooney-Rivlin [40, 45], and Demiray [11] models and their transversely isotropic counterparts, the Lanir [27], Weiss [60], Grooves [15], and Holzapfel [17] models. By design, our neural networks can *interpolate, extrapolate, and explain* the constitutive behavior just like these individual models, and moreover as all these models combined.

### Limitations

Although we have successfully demonstrated the potential of our proposed approach, we see three major limitations: First, and most obviously, the fit of the discovered model and parameters is not perfect and clearly not as good as a fit of first generation Constitutive Artificial Neural Networks or Neural Ordinary Differential Equations. This perceived shortcoming of our method is a result of a combination of two factors, its much lower number of degrees of freedom, and its inherent objective of identifying models that are not just a combination of hyperbolic tangent functions, but rather a generalization of existing computational models. We could address this limitation by adding more terms that still satisfy the polyconvexity condition, for example, more polynomial terms of the invariants or more polyconvex activation functions. However, the downside of this strategy is the loss of a parsimonious representation with interpretable parameters. Second, and this is easily addressed, our current network architecture is limited to discovering constitutive models in which the invariants are fully decoupled. This implies that we cannot discover dispersion-type models in which the individual invariants interact with one another. This limitation is intended to make it easier to account for polyconvexity, but can be easily addressed by a fully connected network architecture in which all nodes of the two hidden layers are interconnected. Third, and probably most difficult to handle, our method remains slightly sensitive to its initial conditions, particularly to the initialization of the network weights. Specifically, even when trained with all available data combined, it occasionally discovers a quadratic term in the second invariant or a quadratic exponential term in the fifth invariant. We tried to regularize the loss function, but neither L1 nor L2 regularization completely eliminates this uniqueness problem. We conclude that this is not an artifact of our method, but rather an indication for the existence of secondary models to explain the data. A crucial next step to gain quantitative insight into all three limitations would be to embed our method into a Bayesian approach to identify the type of outliers in terms of both models and model parameters.

## 7 Conclusion

Constitutive modeling and parameter identification are the cornerstones of continuum mechanics. For decades, the usual standard in constitutive modeling was to first choose a model and then fit its parameters to data. However, this approach depends largely on user experience and personal preference. Here we proposed a new method to simultaneously and fully autonomously discover the best model and parameters to explain experimental data. As a by-product, the method also discovers the best set of experiments to train itself. This is clearly a non-trivial task that, in mathematical terms, translates into a complex non-convex optimization problem. Our solution strategy is to leverage the success, robustness, and stability of the powerful optimization schemes developed for classical neural networks. We formulated the model finding problem as a Constitutive Artificial Neural Network with activation functions representing traditional constitutive models and parameters that have a clear physical interpretation. We have demonstrated the potential of our method for biaxial extension experiments on skin, showing that the network autonomously discovers a small set of non-zero weights that define the discovered model, while the majority of the weights are trained to zero. In contrast to classical neural network modeling, our non-zero weights have a clear physical interpretation and can be translated into well-defined engineering parameters and microstructural features. Our findings suggest that Constitutive Artificial Neural Networks have the potential to enable automated model, parameter, and experiment discovery and could induce a paradigm shift in constitutive modeling, from user-defined to automated model selection and parameterization.

## Data availability

Our source code, data, and examples will be available at https://github.com/LivingMatterLab/CANNs.

## Acknowledgments

This work was supported by a DAAD Fellowship to Kevin Linka and by the Stanford School of Engineering Covid-19 Research and Assistance Fund and Stanford Bio-X IIP seed grant to Ellen Kuhl.

